# FX-Cell: a method for single-cell RNA sequencing on difficult-to-digest and cryopreserved plant samples

**DOI:** 10.1101/2025.03.04.641200

**Authors:** Xin Ming, Mu-Chun Wan, Zheng-Da Zhang, Hao-Chen Xue, Ya-Qi Wu, Zhou-Geng Xu, Yan-Xia Mai, Ying-Xiong Hu, Ke Liu, Jian Gao, Qiao-Lin Shao, D. Blaine Marchant, Brad Nelms, Virginia Walbot, Jia-Wei Wang

## Abstract

Single-cell RNA sequencing (scRNA-seq) in plants requires the isolation of high-quality protoplasts— cells devoid of cell walls. However, many plant tissues and organs are resistant to enzymatic digestion, posing a significant barrier to advancing single-cell multi-omics research in plants. Furthermore, for many field-grown crops, the lack of immediate laboratory access presents another major challenge for protoplast preparation. To address these limitations, we developed the FX-Cell method and its derivatives, FXcryo-Cell and cryoFX-Cell, to enable scRNA-seq with both difficult-to-digest and cryopreserved plant samples. By optimizing the fixation buffer and minimizing RNA degradation, our approach ensures efficient cell wall digestion at high temperatures while maintaining high-quality single cells, even after long-term storage at −80°C, and circumvents use of nuclei, which are not representative of the pool of translatable mRNAs. Using these methods, we successfully constructed high-quality cell atlases for rice tiller nodes, rhizomes of wild rice, and maize crown roots grown in field conditions. Moreover, these methods enable the accurate reconstruction of plant acute wounding responses at single-cell resolution. Collectively, these advancements expand the applicability of plant single-cell genomics across a wider range of species and tissues, paving the way for comprehensive Plant Cell Atlases for plant species.

## Main

The cell holds the genetic blueprint of an organism, yet neighboring cells can differ dramatically in morphology and function. Understanding the gene expression patterns that drive these differences can provide critical insights into the roles, developmental trajectories, and evolutionary histories of cell types, tissues, and entire organisms. Single-cell RNA sequencing (scRNA-seq) has revolutionized our understanding of animal cells, leading to major breakthroughs in cell biology, medicine, and evolution.^1–4^ Applying similar technologies to plants presents significant challenges that hinder the construction of plant cell atlases, the mapping of developmental trajectories, and the functional exploration of genes.^5–11^

The first major challenge arises from the plant cell wall, a rigid outer layer surrounding the plasma membrane. Composed primarily of cellulose, along with other polysaccharides, proteins, and sometimes lignin, the cell wall provides structural integrity but also makes enzymatic digestion difficult. In many plants, secondary cell wall thickening and specialized cell wall components further impede protoplast isolation—the process of releasing live single cells without cell walls.^12–14^

The second limitation is the outdoor cultivation of most plants, especially crops. scRNA-seq requires immediate enzymatic digestion to remove cell walls, yet most farms and field sites lack the necessary molecular biology facilities. As a result, current single-cell transcriptional atlases of crops are derived exclusively from greenhouse-grown materials, which may not adequately capture transcriptome responses under natural conditions.

The third limitation is the time-sensitive nature of plant responses to environmental changes, such as wounding or pathogen infection. Existing protoplast isolation methods require 1-2 hours of enzymatic digestion, making them unsuitable for capturing rapid transcriptional dynamics. A fourth challenge is that while single-nucleus RNA sequencing (snRNA-seq) offers an alternative approach,^15–18^ it often produces lower-quality data than scRNA-seq.^12,19–21^ Additionally, nuclear RNA may not fully represent the cellular transcriptome, in particular the subset of mRNAs that are likely to be translated into protein. Furthermore, isolated nuclei are prone to aggregation, leading to higher doublet rates. Certain transcripts also exhibit differential enrichment between snRNA-seq and scRNA-seq datasets.^22^

To tackle these challenges, we refined existing protoplast isolation protocols and developed simple, versatile methods for scRNA-seq that enable its application to difficult-to-digest and cryopreserved plant samples, as well as samples subjected to wounding treatment. These advancements will expand the range of species and tissues amenable to scRNA-seq, accelerating the development of a comprehensive Plant Cell Atlases,^23,24^ similar to the Human Cell Atlas.^25^

## Results

### Fixation increases single-cell release from plant tissues

Protoplast isolation is perhaps the greatest technical hurdle in scRNA-seq of plant tissues. We hypothesized that the use of coagulant fixatives, which stabilize cells by coagulating the protein matrix while disrupting lipid membranes, followed by cell wall digestion, could offer two key advantages for cell release: fixation (i) stabilizes the cell cytoplasm so that cells can withstand harsher shear forces without breaking, and (ii) allows enzymatic digestion to occur at higher temperatures (∼50°C) where cellulase enzymes are most active.^26^ Additionally, acid treatment can be used and may further degrade cell walls, as acids are commonly used in industrial hydrolysis.^27^

To test this hypothesis, we fixed maize anthers in ice-cold Farmer’s solution (3:1 100% ethanol:glacial acetic acid) and quantified cell release from both fresh and fixed maize anthers. Because fixation disrupts all cellular activities, we refer to isolated structures as “cells” rather than “protoplasts”. We found that an optimized maize anther protoplasting protocol (1.25% w/v Cellulase, 0.5% Pectolyase, 0.5% Macerozyme, 0.5% Hemicellulose)^28^ had a mean release of 4,387 protoplasts per anther after 90 minutes digestion and 11,333 protoplasts per anther when extended to 16 hours (Fig. 1a). In comparison, if anthers were fixed prior to digestion in a reduced enzyme mix (1.25% w/v Cellulase and 0.4% Macerozyme), 15,900 cells were released within 90 minutes; this increase in cell release is presumably because the cells were stabilized against mechanical lysis by fixation, while unfixed cells are very fragile. When incubation temperature was increased from 30°C (standard) to 50°C we observed an average release of 45,033 cells (Fig. 1a), close to the actual number of 50,000 cells in a 2.0 mm maize anther.^29^

**Fig. 1.**
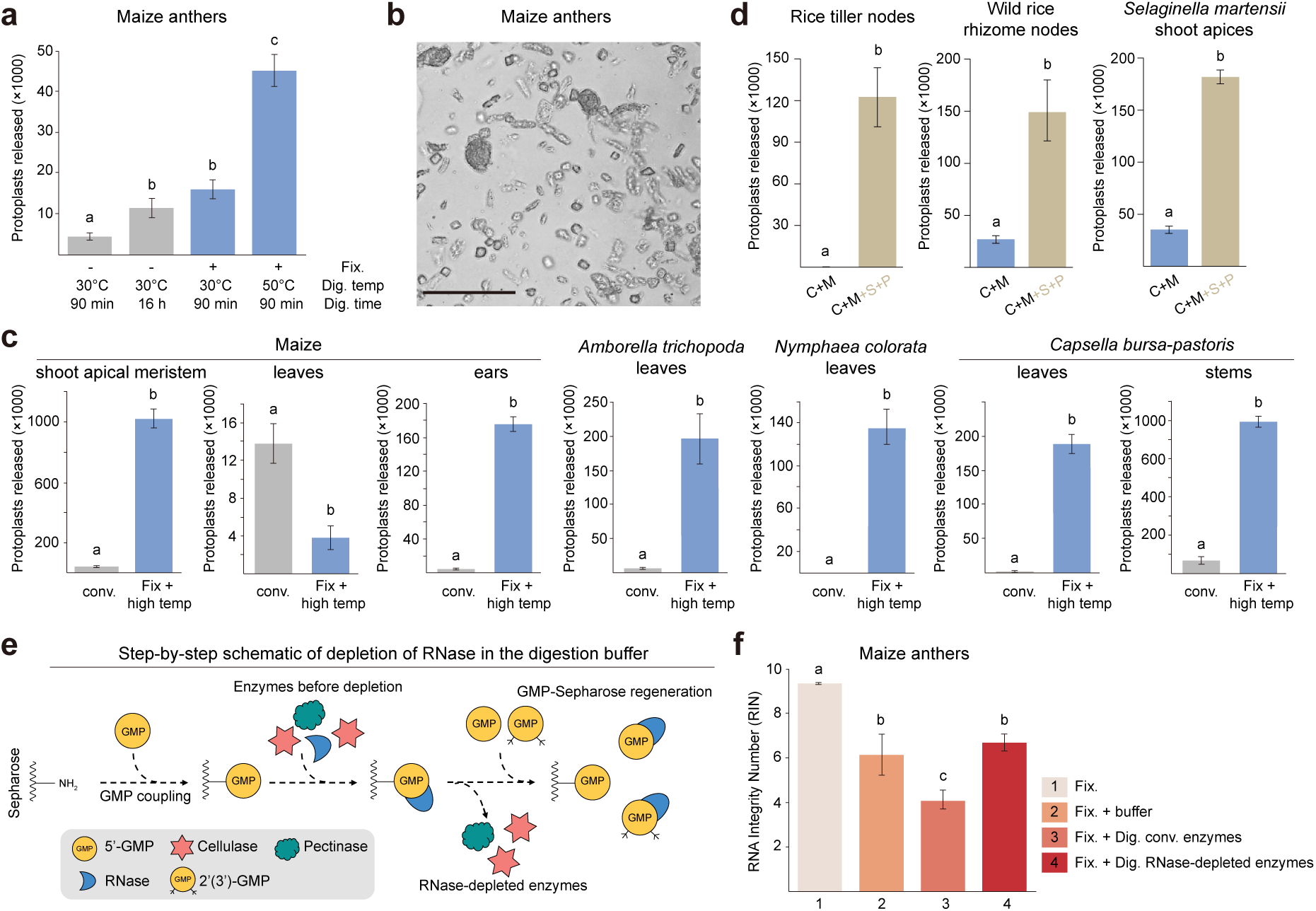
Fixation and digestion at high temperatures protocol facilitates cell release. **a,** Quantification of released protoplasts or cells from maize anthers using different protocols. Maize anthers were digested for 90 minutes or 16 hours at 30°C (grey bars)^28^ with optimized enzyme mix (1.25% w/v Cellulase-RS, 0.5% Pectolyase Y-23, 0.5% Macerozyme-R10, 0.5% Hemicellulose) or first fixed and then digested for 90 minutes with reduced enzyme mix (1.25% w/v Cellulase-RS and 0.4% w/v Macerozyme-R10) at either 30°C or 50°C (blue bars). The number of released protoplasts or cells was quantified using a hemocytometer. Data represent the mean ± s.e.m. from five independent experiments. Different letters indicate statistically significant differences (Student’s *t*-test, *p* < 0.05). **b,** Representative image of released cells from fixed maize anthers. Scale bar, 100 μm. **c,** Quantification of released protoplasts or cells from different tissues and species. Plant tissues were subjected to either fresh protoplasting or fixation and digestion at high temperatures as described in (**a**). Data represent the mean ± s.e.m. from five independent experiments. **d,** Quantification of released cells from fixed rice tiller nodes, wild rice rhizome nodes, and shoot apices of *Selaginella martensii*. Tissues were enzymatically digested at a high temperature using two enzymes (i.e., reduced enzyme mix, C+M, 1.25% w/v Cellulase-RS and 0.4% w/v Macerozyme-R10) or four enzymes (C+M+S+P, 1% w/v Cellulase-RS, 0.5% w/v Macerozyme-R10, 1% w/v Snailase, and 0.5% w/v Pectinase). Data represent the mean ± s.e.m. from three independent experiments. **e,** Schematic representation of RNase depletion in the digestion buffer. 5’-GMP was coupled onto NH_2_-Sepharose and used to remove RNase from the digestion enzymes. Following purification, the GMP-Sepharose column was regenerated by adding 5’-GMP and 2’(3’)-GMP (referred to tri-GMP). **f,** RNA quality assessment of fixed maize anthers under different conditions: (1) without digestion, (2) digested at 50°C for 90 minutes in enzyme buffer lacking enzymes, (3) digested at 50°C for 90 minutes using commercial enzymes, and (4) digested at 50°C for 90 minutes using RNase-depleted enzymes. Data represent the mean ± s.e.m. from five independent experiments.

The optimized maize anther cell release protocol with unfixed tissue yielded fewer than expected epidermal and endothecial cells, both of which tended to remain clumped and undigested, producing a skewed release favoring tapetal cells, middle layer cells, and meiocytes. When the anthers were fixed then digested at 50°C we did not observe any cell clumps, debris, or undigested material, suggesting that the digestion was complete (Fig. 1b). In addition to increasing the cell release efficiency and cell type representation, we found that fixation prior to digestion maintained cells’ natural morphology allowing the potential for cell type identification post-isolation (Fig. 1b; Extended Data Figs. 1a,b).

To assess the method’s broader applicability, we tested four additional maize tissues (shoot apical meristem, leaf, root, and young ear) and three non-model plant taxa (*Amborella trichopoda* leaf, *Nymphaea colorata* leaf, and *Capsella bursa-pastoris* leaf and stem) (Fig. 1c; Extended Data Figs. 1a,c). As shown in Extended Data Fig. 1a, cellular morphology was maintained in each fixed sample, allowing obvious differentiation of the varying cell types. Notably, cell release was 10- to 364-fold higher in fixed tissues compared to a fresh sample, with the exception of maize leaves in which there were 3.6 as many cells released from a fresh sample compared to the fixation-based protocol (Fig. 1c). Presumably, cells with large fluid filled vacuoles, such as maize mesophyll, are very fragile after fixation, because they have too few coagulated proteins.

Specialized cell wall components in some plants necessitated modifications to the digestion buffer. The fixation-digestion at high temperatures protocol using reduced enzyme mix failed to efficiently digest rice tiller nodes, wild rice (*Oryza longistaminata*) rhizome nodes, and lycophyte (*Selaginella martensii*) shoot apices. To address this, we incorporated 1% Cellulase, 1% Snailase, 0.5% Macerozyme, and 0.5% Pectinase, which significantly improved digestion by targeting chitin and pectin (Fig. 1d; Extended Data Fig. 1b). Overall, our fixation-based protocol, combined with an expanded enzyme mix, effectively dissociated diverse plant tissues into single cells, broadening the applicability of scRNA-seq to previously challenging species and tissues.

### RNase-depletion of the digestion enzyme cocktail is necessary for maintaining RNA quality

While fixation itself does not affect RNA quality, it removes the cell membrane and makes the internal RNA contents accessible to RNases in solution. This generates a challenge during enzymatic digestion, because most cell wall digesting enzymes are complex mixtures that contain substantial RNase activity. We tested several RNase inhibitors, including commercial inhibitors, EDTA, and vanadyl ribonucleoside complexes, but found none that could effectively inhibit the RNase activity in cell wall digestion enzyme blends. This is partly because many available RNase inhibitors target the RNase A family of enzymes,^30^ which is only produced in vertebrates. Secreted fungal RNases are primarily of the T1 and T2 families.^30^

To surmount this complication, we adapted a column-based method to reduce fungal T1 and T2 RNases by binding them to Sepharose/Agarose coupled with guanosine monophosphate (GMP) (Fig. 1e).^31^ We found that cell wall digesting enzymes readily passed through GMP-Sepharose/Agarose columns, while the contaminating RNases remained bound. After column depletion, RNase activity was greatly reduced in the digestion enzyme blend (Extended Data Fig. 1d). The RNase-depleted enzymes remained stable when stored as glycerol stocks for at least one year at −20°C.

Because GMP-Sepharose/Agarose synthesis involves multiple steps and lacks standardized quality control, we implemented an assessment step to verify successful GMP coupling. Specifically, GMP-Sepharose/Agarose was incubated in hydrochloric acid in a boiling water bath for 1 hour, and its optical density at 248 nm was measured (Extended Data Fig. 1e). Additionally, we introduced a regeneration step using a wash buffer containing 5 M NaCl and 1 mM tri-GMP (a mixture of 1 mM 5’-GMP and 1 mM 2’(3’)-GMP) to remove bound RNases, allowing column reuse (Fig. 1e; see Methods). These modifications enable laboratories with limited biochemistry expertise to easily purify digestive enzymes of RNases.

We next tested the effect of the fixed tissue dissociation procedure on RNA quality. RNA isolated from fixed maize anthers had an average RNA Integrity Number (RIN) of 9.3 demonstrating fixation did not cause any significant decrease in RNA quality (Fig. 1f). Fixed anthers digested at 50°C in a commercial enzyme blend had a RIN of 4.1 with very noticeable loss of ribosomal RNA. After fixation then digestion with RNAse-depleted enzymes, the RIN was 6.7 demonstrating the fixed tissue dissociation protocol can produce RNA of reasonable quality, although there is a decrease in RNA integrity relative to undigested tissue. When fixed anthers were incubated in enzyme buffer at 50℃ without enzymes, we observed a similar RIN of 6.1. Therefore, the decrease in RNA integrity during incubation is not exogenous enzyme-dependent, rather we suspect this degradation is caused by endogenous anther RNases that survive the fixation process.

### Development of the FX-Cell method

To evaluate the applicability of the fixation-digestion at high temperatures method for scRNA-seq, maize anthers were fixed and digested with RNAse-depleted enzymes at 50°C for 90 minutes. The resultant cells were sorted and isolated using either BioSorter (Union Biometrica) or Hana (Namocell) machines into 96-well plates, and scRNA-seq libraries were prepared using a modified CEL-Seq2 library preparation protocol.^28,32^ Of the 384 maize anther single cells examined, 307 exhibited more than 500 unique molecular identifiers (UMIs) and at least 200 detected genes after the removal of cell-cycle genes. An average of 5,885 UMIs and 2,016 transcribed genes per cell were detected (Extended Data Table 1). The dataset was classified into four distinct clusters, two of which were reclustered based on marker gene expression, producing six total clusters (Extended Data Table 1). Notably, clusters corresponding to meiocyte, endothecium, tapetum, and epidermis were identified (Extended Data Fig. 2). A more extensive dataset generated using this method enabled the reconstruction of developmental trajectories of all living maize anther cell types, spanning from early cell proliferation to subsequent cell fate determination and differentiation over a 20-day time course.^33^

Encouraged by these findings, efforts were made to determine whether this method could be adapted for commercial high-throughput scRNA-seq platforms such as Chromium (10× Genomics). To facilitate direct comparisons with conventional scRNA-seq and snRNA-seq, rice root tissues with existing scRNA-seq data were selected as the experimental samples.^34^ The data quality obtained using the fixation and digestion at high temperatures method was suboptimal. While the median number of transcribed genes detected per cell was 1,230 and 1,369 (Fig. 2b; Extended Data Table 1), the total number of captured cells was only 2,084 and 2,810, respectively (Fig. 2a; Extended Data Table 1). The cumulative cell count across two biological replicates was significantly lower than the 27,469 cells obtained from traditional protoplast-based scRNA-seq,^34^ indicating a considerable number of low-quality cells were generated, leading to their automatic removal by Cell Ranger. Consistently, when data quality (i.e., the number of genes per cell) was not considered, the filtering criteria required to retain 8,000 cells resulted in a decrease in the median number of transcribed genes per cell, dropping to 303 and 444 (Fig. 2c; Extended Data Table 1). The data quality for snRNA-seq was similarly unsatisfactory, with single-cell gene counts of 635 and 629, and captured cell numbers of 1,428 and 1,835 (Figs. 2a,b; Extended Data Table 1).

**Fig. 2.**
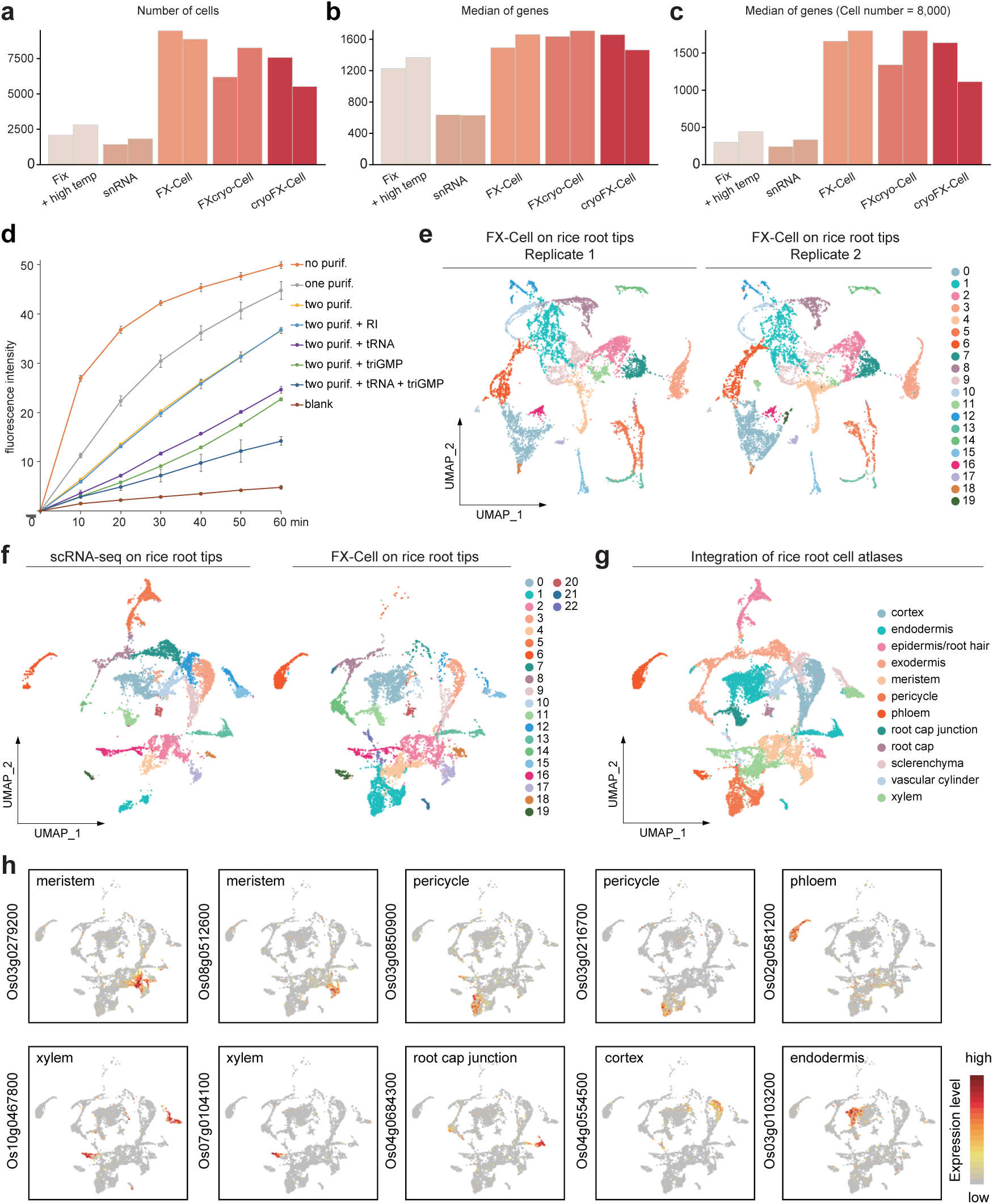
Development and evaluation of FX-Cell for high-throughput scRNA-seq. **a,b,** The number of recovered cells (**a**) and the median number of transcribed genes (**b**) in the rice root tip cell atlas generated by different scRNA-seq methods. Two biological replicates were performed for each method. **c,** The median number of transcribed genes in the rice root tip cell atlas under conditions where filtering criteria were adjusted to achieve 8,000 cells in the atlas, without considering data quality (i.e., the number of genes per single cell). The same data as in (**a,b**) were used. **d,** Assessment of RNase enzymatic activity in the digestion solution under various conditions. “purif” indicates RNase removal via affinity chromatography. “RI” refers to a commercial RNase inhibitor. “triGMP” represents a mixture of 5’-GMP and 2’(3’)-GMP. “Blank” serves as a control using the enzyme-dissolving RBB buffer. Data represent the mean ± s.e.m. from three independent experiments. **e,** UMAP plots depicting the rice root tip cell atlas generated by FX-Cell. Two biological replicates were performed. The batch effect was not corrected using the Harmony algorithm. Different colors indicate distinct cell clusters. **f,** Comparison of rice root tip cell atlases generated by FX-Cell and traditional protoplasting-based scRNA-seq.^34^ The datasets were integrated, and batch effects were removed using the Harmony algorithm. Different colors represent distinct cell clusters. To minimize biases due to differences in cell numbers across datasets, both atlases were normalized to approximately 9,000 cells. **g,** Annotation of the integrated rice root tip cell atlas shown in (**f**) based on published cell type annotation.^34^ Different colors correspond to distinct cell types. **h,** UMAP plots illustrating the expression patterns of known cell type marker genes in the rice root tip cell atlas generated by FX-Cell, corresponding to (**f**).

These results likely stem from RNA degradation, prompting efforts to improve RNA integrity and increase the number of transcribed genes captured per cell. During the digestion of rice root tips, elevated temperatures and prolonged digestion times enhanced RNA degradation and decreased RNA integrity (Extended Data Figs. 1f-i). To address this issue, digestion time was reduced from 90 minutes to 30 minutes, and the temperature was lowered from 50°C to 40°C. Additionally, to minimize RNase contamination in the digestion buffer, digestion enzymes were purified twice using GMP-Sepharose columns (Fig. 2d). Remarkably, the incorporation of tRNA and tri-GMP effectively inhibited RNase activity (Fig. 2d).

Following these optimizations, the experiment was repeated using rice root tips with two biological replicates. Compared to the original method and the data obtained from snRNA-seq, the optimized protocol resulted in a substantial increase in the number of detected cells (9,474 and 8,874 cells) as well as in the median number of transcribed genes per cell (1,494 and 1,661 genes) (Figs. 2a,b; Extended Data Table 1). Notably, when the filtering criteria were adjusted to retain 8,000 cells, the gene counts per individual cell increased to 1,661 and 1,801, which were substantially higher than those obtained using the original method and snRNA-seq (Fig. 2c; Extended Data Table 1). Furthermore, Uniform Manifold Approximation and Projection (UMAP) visualization demonstrated high reproducibility between the two independent experiments (Fig. 2e). The dataset generated using the optimized method also displayed high consistency with traditional protoplasts-based scRNA-seq data (Fig. 2f).^34^ Using published cell type annotation,^34^ most of the cell types present in rice root tips were successfully identified (Fig. 2g). Importantly, the expression specificity of these marker genes was largely preserved (Fig. 2h; Extended Data Fig. 3b). The lower number of root hair cluster cells in the new atlas may be attributed to developmental differences arising from variations in cultivation conditions (Extended Data Fig. 3c).

In summary, a novel high-throughput plant scRNA-seq method, termed FX-Cell, was developed. This method outperforms snRNA-seq and produces data of comparable quality to that obtained using traditional scRNA-seq. Importantly, FX-Cell enhances the efficiency of single-cell release, enabling the application of scRNA-seq to tissues and organs that are otherwise challenging to digest and enhances the likelihood of recovering and analyzing rare cell types.

### Development of FXcryo-Cell and cryoFX-Cell for cryopreserved plant samples

Traditional protoplast-based scRNA-seq methods require the use of fresh plant tissues, necessitating immediate processing after sampling, which significantly limits their applicability. The ability to perform scRNA-seq on fixed plant materials following cryopreservation would greatly expand the potential applications of this technology. To test this hypothesis, fixed rice roots were cryopreserved by rapidly freezing them in liquid nitrogen and then stored at −80°C in an ultra-low temperature freezer. After thawing and subsequent cell wall digestion at 40°C for 30 minutes, the morphology of the cells released from cryopreserved rice roots was largely similar to that of non-cryopreserved roots (Fig. 3c). Similar observations were made in other plant tissues, including *A. thaliana* leaves and *S. martensii* shoot apices (Extended Data Figs. 4a,b).

**Fig. 3.**
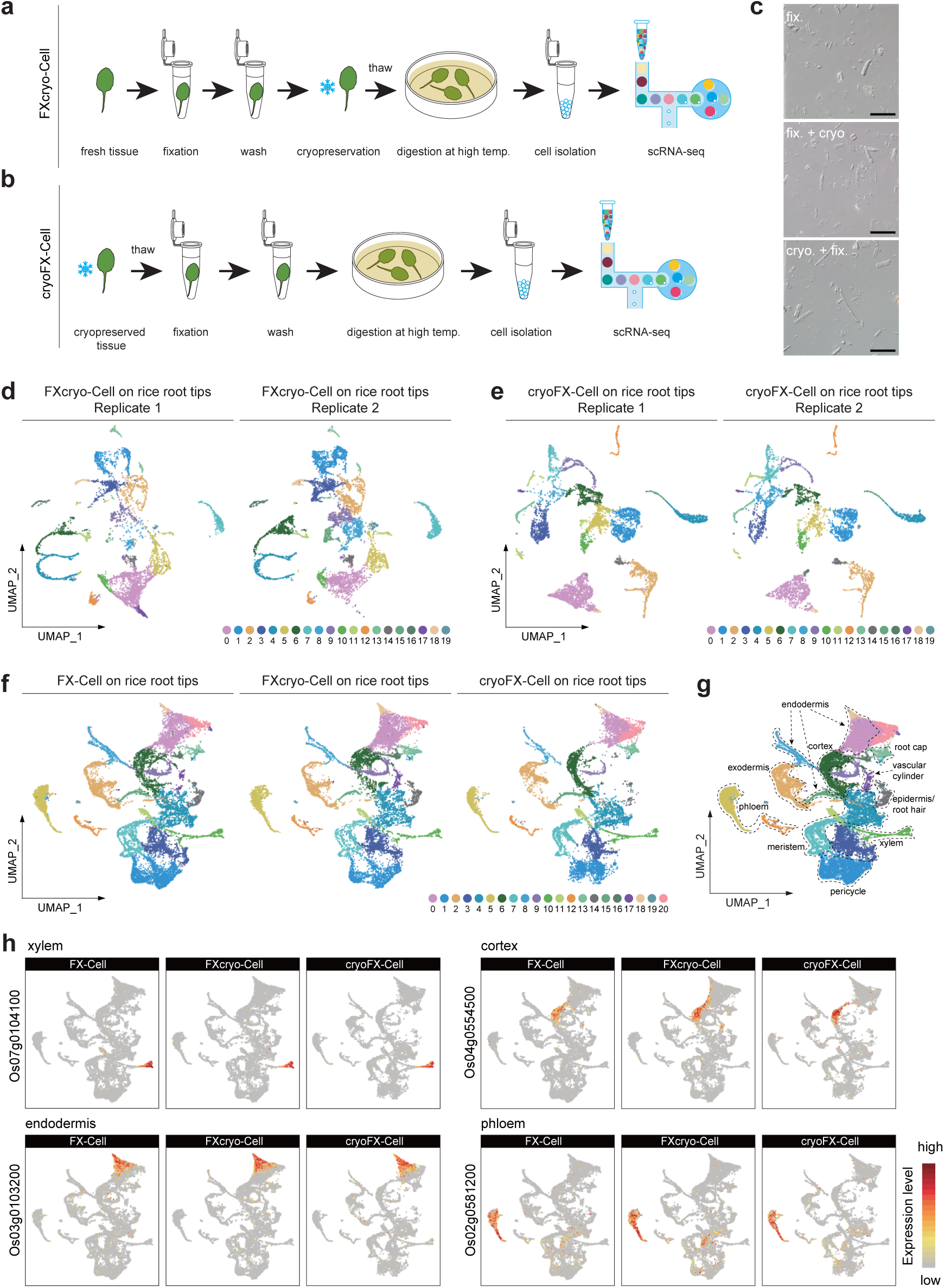
Applications of FXcryo-Cell and cryoFX-Cell on rice root tip samples. **a,b,** Step-by-step schematic of the FXcryo-Cell (**a**) and cryoFX-Cell (**b**) workflows. **c,** Representative images showing released cells from rice root tips. Cells were obtained from tissues subjected to three different treatments: (1) fixation followed by high-temperature enzymatic digestion (fix.), (2) fixation, freezing, thawing, and then high-temperature enzymatic digestion (fix. + cryo.), and (3) freezing, thawing, fixation, and high-temperature enzymatic digestion (cryo. + fix.). Scale bars, 50 μm. **d,e,** UMAP plots displaying rice root tip cell atlases generated using FXcryo-Cell (**d**) and cryoFX-Cell (**e**). Two biological replicates were performed for each experiment. Batch effects were not removed by the Harmony algorithm. Different colors indicate distinct cell clusters. **f,** Joint analysis of rice root tip cell atlases from FX-Cell, FXcryo-Cell, and cryoFX-Cell. Batch effects were not removed by the Harmony algorithm. Different colors represent different cell clusters. **g,** Integrated rice root tip cell atlas, annotated based on published cell type marker genes (Extended Data Fig. 3a; Extended Data Table 1). Atlases from FX-Cell, FXcryo-Cell, and cryoFX-Cell were combined. Different colors represent different cell clusters, with related clusters grouped by dashed circles. **h,** UMAP plots showing expression patterns of representative cell type marker genes in the integrated rice root tip cell atlas. Four representative cell types are displayed. See Extended Data Fig. 4d for additional cell types.

scRNA-seq was then performed on fixed and cryopreserved rice root tips using two biological replicates, yielding median gene counts of 1,637 and 1,711 per individual cell and capturing 6,192 and 8,259 cells, respectively (Figs. 2a,b; Extended Data Table 1). When filtering criteria were adjusted to retain 8,000 cells, the median gene counts per individual cell reached 1,341 and 1,802, surpassing the results obtained from snRNA-seq (Fig. 2c; Extended Data Table 1). UMAP visualization demonstrated distinct cell clustering and high reproducibility between the two independent experiments (Fig. 3d). This method was therefore designated as FXcryo-Cell (Fig. 3a).

Under certain circumstances, such as field sampling, immediate fixation followed by freezing can be challenging. To address this, an alternative approach was developed in which plant materials were directly cryopreserved and fixed immediately after thawing, followed by high-temperature enzymatic digestion. The quality of cells obtained using this workflow was comparable to that of FXcryo-Cell (Fig. 3c; Extended Data Figs. 4a-c). Notably, scRNA-seq performed on these cells from rice root tips, using two biological replicates, yielded high-quality and reproducible data (Fig. 3e), with median gene counts per individual cell of 1,660 and 1,465 and total captured cell numbers of 7,578 and 5,524 cells, respectively (Figs. 2a,b; Extended Data Table 1). When 8,000 cells were retained in Seurat analysis, the median gene counts per cell were 1,640 and 1,115 (Fig. 2c; Extended Data Table 1). This method was designated as cryoFX-Cell (Fig. 3b).

To evaluate the consistency among FXcryo-Cell, cryoFX-Cell, and FX-Cell, datasets from rice roots generated by these three methods were merged without batch effect correction. The cell types were annotated with published cell type marker genes (Fig. 3g; Extended Data Fig. 3a; Extended Data Table 1). UMAP visualization demonstrated strong reproducibility across all three datasets (Figs. 3f,g). Additionally, marker genes specifically expressed in previously published scRNA-seq data of rice roots were also specifically expressed in both FXcryo-Cell and cryoFX-Cell datasets (Fig. 3h; Extended Data Fig. 4d).

In conclusion, FXcryo-Cell and cryoFX-Cell effectively replace traditional scRNA-seq methods, significantly expanding the potential applications of scRNA-seq in plants. In the following sections, three potential application scenarios for these two methods are explored.

### Application of FXcryo-Cell on difficult-to-digest plant samples

The shoot architecture in cereals is largely influenced by tillers, which are lateral shoots arising from axillary buds at basal nodes (insets in Fig. 4a).^35^ Dissecting the development of basal tiller nodes at single-cell resolution is therefore crucial for understanding crop architecture and yield. As previously mentioned, however, traditional enzymatic digestion methods struggle to obtain protoplasts from these structures. FXcryo-Cell was found to effectively remove cell walls from basal tiller nodes of 5-day-old rice plants, releasing a greater number of individual cells (Fig. 1d). Consequently, an atlas of 5,822 high-quality cells was obtained, with a median transcribed gene count of 1,504 (Extended Data Table 1). Established marker genes were used to annotate cell clusters corresponding to various cell types (Fig. 4a; Extended Data Fig. 5a; Extended Data Table 1). A central meristem-like cell cluster (cluster 0) was surrounded by differentiating cell types, including phloem (clusters 8 and 9), xylem (cluster 10), cambium (cluster 13), endodermis (clusters 1 and 3), and epidermis (cluster 5). Additionally, consistent with morphological observations indicating that both root and tiller bud formation occurs at the basal tiller node (insets in Fig. 4a), a root cap cell cluster (cluster 11) was identified.

**Fig. 4.**
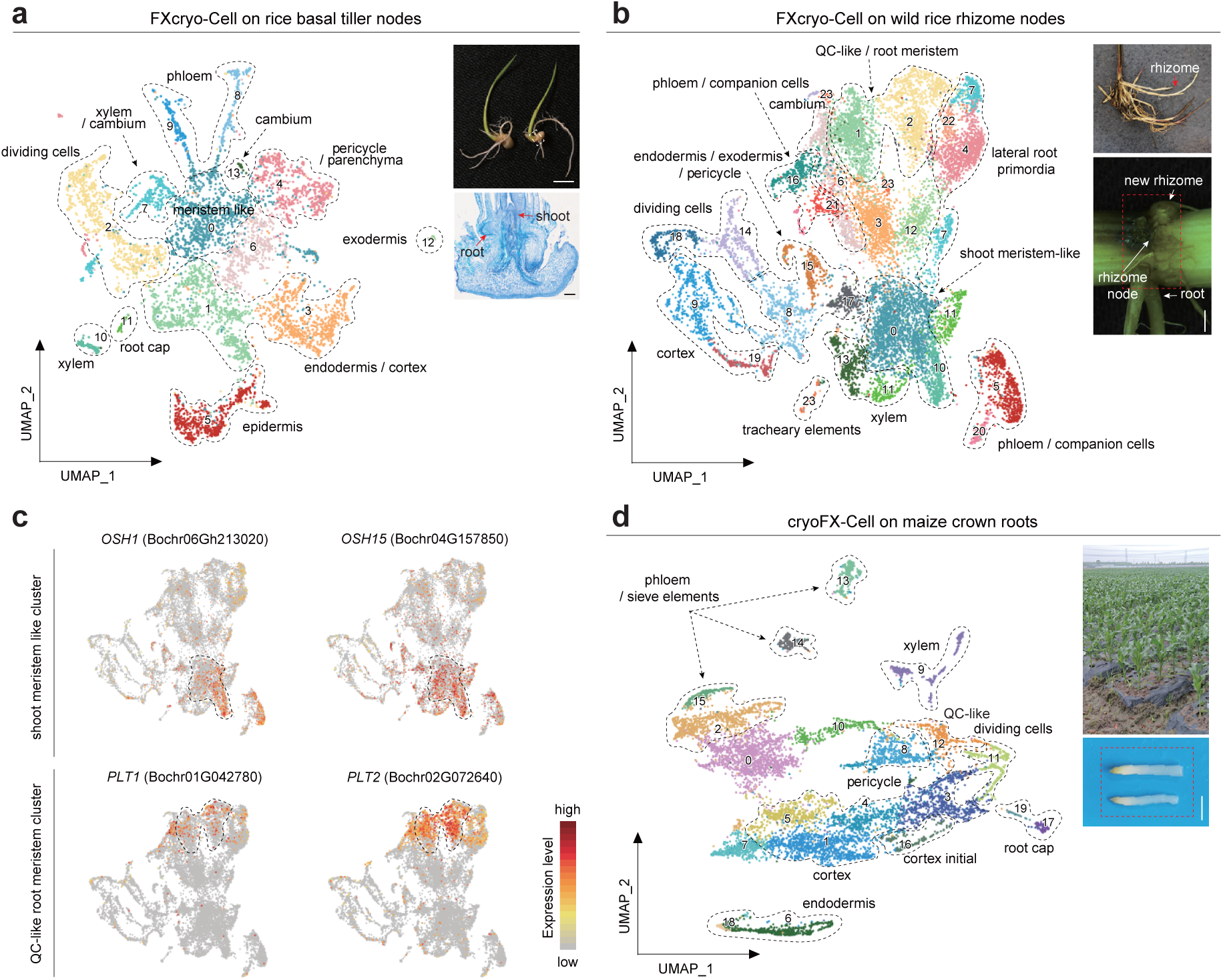
Cell atlases of difficult-to-digest and cryopreserved plant samples. **a,** UMAP plot of the cultivated rice basal tiller node cell atlas generated using FXcryo-Cell. Different colors indicate distinct cell clusters, with related clusters grouped by dashed circles. Insets: rice seedlings (upper panel) and a longitudinal section of a basal tiller node (lower panel). The harvested tissue used for FXcryo-Cell is highlighted by a white dashed box. The basal node can give rise to both tiller buds and roots (red arrows). The atlas was annotated based on published cell type marker genes (Extended Data Fig. 5a; Extended Data Table 1). Scale bars: 0.5 cm (upper panel), 200 μm (lower panel). **b,** UMAP plot of the wild rice rhizome node cell atlas generated using FXcryo-Cell. Insets: growing rhizomes (upper panel, red arrow) and the harvested rhizome node (red dashed box) used for FXcryo-Cell. The atlas was annotated based on published cell type marker genes (Extended Data Fig. 5b; Extended Data Table 1). Scale bar, 2 mm. **c,** UMAP plots showing the expression patterns of shoot and root meristem identity genes in the wild rice rhizome node cell atlas (**b**). QC-like/root meristem and shoot meristem-like cell clusters are circled. **d,** UMAP plot of the cell atlas of field-grown maize emerging crown roots generated using cryoFX-Cell. Insets: field-grown maize plants (upper panel) and the harvested crown roots (red dashed box) used for cryoFX-Cell. The atlas was annotated based on published cell type marker genes (Extended Data Fig. 6a; Extended Data Table 1). Scale bar, 0.5 cm.

To further validate the applicability of FXcryo-Cell for difficult-to-digest plant samples, rhizomes— underground, horizontal stems that facilitate the perennial growth habit in wild rice (*O. longistaminata*) (insets in Fig. 4b)—were analyzed.^36–38^ Similar to tiller nodes, rhizome nodes could not be effectively digested into protoplasts using traditional digestion methods. FXcryo-Cell successfully overcame this limitation (Fig. 1d). As shown in Fig. 4b, the application of FXcryo-Cell on rhizome nodes at the budding stage generated a high-quality single-cell resolution map, capturing 9,233 cells with a median gene count of 891 per cell (Extended Data Table 1). The majority of cell types were characterized using published rice and *A. thaliana* cell type marker genes (Fig. 4b; Extended Data Fig. 5b; Extended Data Table 1).^34,39,40^

New buds on rhizomes either grow underground as new rhizomes or emerge aboveground as aerial shoots.^37^ Accordingly, shoot meristem-like clusters (clusters 0 and 10) were identified in the rhizome node atlas (Fig. 4b). These clusters showed high expression of *ORYZA SATIVA HOMEOBOX1* (*OSH1*) and *OSH15* (Fig. 4c), genes associated with shoot meristem identity.^41,42^ As shown in Fig. 4b, rhizome nodes were observed to generate roots. Correspondingly, quiescent center (QC)-like/root meristem clusters (clusters 1 and 2) were detected (Fig. 4b), with strong expression of homologs of the well-known *A. thaliana* root identity *PLETHORA* (*PLT*) genes, including *PLT1* and *PLT2* (Fig. 4c).^43^

Collectively, these findings demonstrate that FXcryo-Cell is a valuable tool for studying difficult-to-digest plant samples. The improved method can provide fresh insights into the complexity of cell types of both tiller and rhizome nodes in rice.

### Application of cryoFX-Cell to field samples

Compared to greenhouse-grown samples, applying scRNA-seq to field-collected plant materials provides a more accurate representation of plant responses to complex and dynamic environmental conditions at the cellular level. As previously mentioned, traditional protoplasting-based scRNA-seq methods require immediate processing of freshly collected plant samples, significantly limiting their use in field-based research. The optimized cryoFX-Cell method enables the use of cryopreserved plant samples for single-cell preparation, offering an ideal solution for field sample analysis.

To evaluate this approach, emerging crown roots were collected from 45-day-old field-grown maize (insets in Fig. 4d). The samples were rapidly frozen in dry ice and stored at −80°C for long-term preservation (Fig. 4d). Following a 50-day storage period, a cryoFX-Cell experiment was conducted. The results yielded 9,127 captured cells, with a median gene count of 1,857 per cell (Extended Data Table 1). Cell clusters were annotated based on established marker genes and published references (Fig. 4d; Extended Data Fig. 6a).^44–46^ Notably, consistent with previous findings that primary and crown roots have similar anatomy,^47^ the vast majority of maize primary root cell types identified using traditional scRNA-seq methods were also recovered (Fig. 4d; Extended Data Figs. 6b,c).^45^

### Application of FXcryo-Cell to probe acute wounding responses at single-cell resolution

As sessile organisms, plants have evolved complex mechanisms to cope with dynamic environmental challenges. In response to wounding, calcium and jasmonic acid signaling pathways are rapidly activated at the site of damage, initiating tissue repair and organ regeneration.^48–51^ To date, studies have primarily relied on bulk samples,^48,52^ leaving it unclear whether different plant cell types exhibit distinct responses to wounding. To address this question, transcriptomic changes at the single-cell level must be examined. Traditional plant scRNA-seq methods require protoplast preparation, which may inadvertently introduce wounding effects into control samples, complicating the acquisition of ground-truth data from unwounded plants. Fixation preserves the original state of plants at the time of sampling, effectively mitigating this issue.

To evaluate FXcryo-Cell in this context, the third-to-last true leaves of 24-day-old *A. thaliana* plants were wounded. Intact (control) and wounded leaves were harvested 2 hours post-injury, fixed, and washed twice with 0.1× PBS, rapidly frozen in liquid nitrogen, and then cryopreserved at −80°C. After five days, the samples underwent FXcryo-Cell processing (Fig. 3a). The intact leaf sample captured 7,815 cells, with an median of 3,746 transcribed genes per cell, while the wounded sample captured 8,979 cells, with an median of 4,012 transcribed genes per cell (Extended Data Table 1).

To compare FXcryo-Cell with traditional protoplasting-based scRNA-seq, previously published enzymatic digestion scRNA-seq data from the Chory Laboratory^53^ were retrieved. Batch effect correction was performed using the Harmony algorithm, integrating these datasets with those generated by FXcryo-Cell. As shown in Fig. 5a, the integrated cell atlas demonstrated strong concordance across the three datasets. Cell clusters were annotated based on established *A. thaliana* leaf scRNA-seq studies and known marker genes (Fig. 5b; Extended Data Fig. 7a; Extended Data Table 1).

**Fig. 5.**
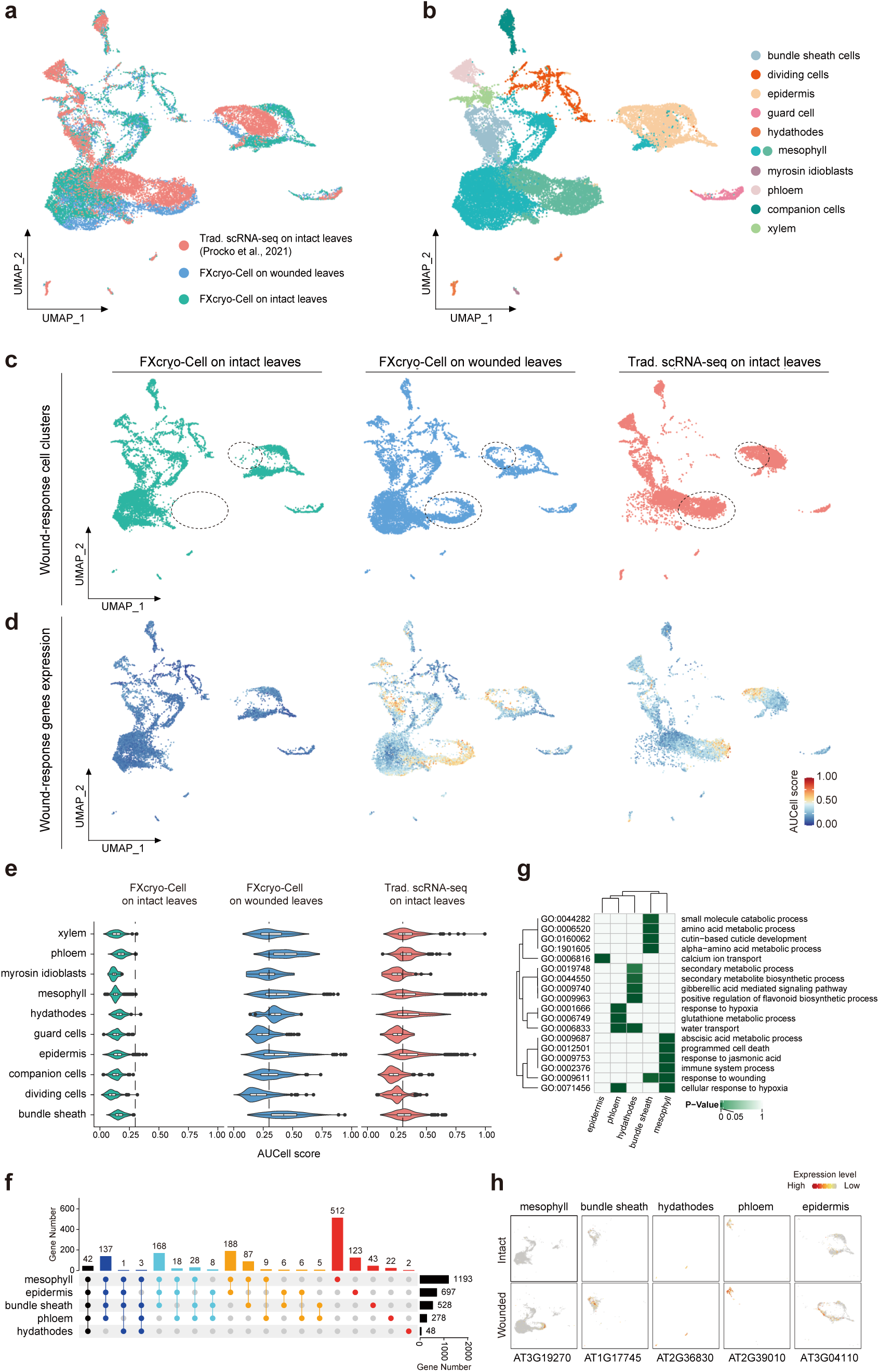
Probing wound response in *A. thaliana* leaves at single-cell resolution using FXcryo-Cell. **a,** Joint analysis of intact (green) and wounded (blue) *A. thaliana* leaf cell atlases generated using FXcryo-Cell, alongside the intact *A. thaliana* leaf cell atlas (red) obtained from traditional protoplasting-based scRNA-seq.^53^ The Harmony algorithm was applied to correct batch effects. Different colors indicate distinct scRNA-seq datasets. **b,** Annotation of the integrated cell atlas from (**a**), based on published cell type marker genes (Extended Data Fig. 7a; Extended Data Table 1). Different colors represent distinct cell types. **c,** Comparative analysis of the three atlases shown in (**a**). Different colors represent distinct scRNA-seq datasets. Wound-responsive epidermal and mesophyll cells are circled. Please note that these cells were barely detected in the intact leaf atlas generated by FXcryo-Cell (green), but evident in that generated by traditional protoplasting-based scRNA-seq (red). **d,** UMAP plots displaying expression levels of wound-response genes across the three datasets from (**a**). Wound-response genes extracted from GO terms (GO: 0009611; Extended Data Table 2) were used as the gene set and analyzed using AUCell. **e,** Wound responsiveness across cell types. Wound-response genes extracted from GO terms (GO: 0009611; Extended Data Table 2) were the gene set. AUCell scores for each cell type were calculated and plotted. **f,** Upset plot presenting shared and unique wound-induced genes in each cell type. The number of genes for each cell type is presented on the top of the colored bars. Total number of wound-induced genes in each cell type is presented on the right. **g,** GO term analysis of unique wound-induced genes in each cell type. **h,** UMAP plots presenting representative cell-type specific wound-induced genes. Only cells corresponding to each specific cell type are presented. The upper and lower rows indicate intact and wounded *A. thaliana* leaf cell atlases produced using FXcryo-Cell (**c**), respectively.

Compared to intact leaves, wounded leaves exhibited two additional cell clusters corresponding to epidermal and mesophyll cells (Fig. 5c). Many wound-induced genes were detected exclusively in wounded leaves (Extended Data Fig. 7b).^52^ Genes associated with plant wounding responses (GO: 0009611, Extended Data Table 2) were extracted and analyzed using AUCell,^54^ confirming that these unique cell populations were strongly linked to wound response pathways (Fig. 5d). Notably, scRNA-seq data obtained using traditional protoplasting exhibited greater similarity to FXcryo-Cell data from wounded leaves (Figs. 5c,d; Extended Data Fig. 7b), suggesting that stress-related gene expression patterns in existing published plant cell atlases may not fully reflect true biological states, potentially including cell clusters that do not naturally exist.

Importantly, comparison atlases generated from intact and wounded leaves using FXcryo-Cell enabled us, for the first time, to investigate wound responses in plants at the single-cell or cell-type levels. Interestingly, we found that different cell types did not respond uniformly to wounding: dividing cells exhibited the lowest wound responses, while epidermal, mesophyll, phloem, and bundle sheath cells displayed a stronger response (Fig. 5e; Extended Data Table 2). Notably, while different cell types had common wound-response genes, they also possessed their own unique wound-response genes (Figs. 5f,h; Extended Data Figs. 8a,c; Extended Data Table 2), with this phenomenon being particularly pronounced in mesophyll cells. GO term analysis indicated that these cell-type-specific wound-response genes may be associated with various biological processes (Fig. 5g; Extended Data Fig. 8b; Extended Data Table 2). For instance, genes in mesophyll cells were enriched for terms linked to abscisic acid metabolic processes, programmed cell death, and response to jasmonic acid. Conversely, epidermal cells were associated with terms linked to calcium ion transport (Fig. 5g). These findings indicate that plant wound perception is more complex than previously envisioned, with different cell types responding to wound signals differently. The biological significance underlying these differences requires further exploration.

Taken together, FXcryo-Cell provides unprecedented advantages for investigating acute plant wounding responses. By preserving the plant’s original state at the time of sampling, this method is particularly beneficial for experiments requiring rapid processing, especially in challenging or time-sensitive sampling conditions. In the future, FXcryo-Cell could be applied to plant-pathogen interactions and responses to diverse abiotic stresses.

## Discussion

In summary, the applications presented above demonstrate that FX-Cell and its derivatives FXcryo-Cell and cryoFX-Cell significantly expand the utility of scRNA-seq in plants. As with any new methods, potential limitations must be considered. The primary advantage of these techniques lies in their ability to enhance the release of single cells from plant tissues. Nonetheless, certain contexts present challenges. First, some plant cells have large fluid filled vacuoles and are very fragile after fixation; for instance, we found maize leaf mesophyll cells do not hold up well to our method. As a result, other approaches may be better for cells with very high water content. With any new tissue, we recommend first testing this method using commercially available enzymes to see how well the cells of interest are successfully released before committing to RNase-depletion of the enzymes.

Second, the large size and irregular morphology of many plant cells (10-100 μm) compared to animal cells (10-30 μm) may lead to clogging in microfluidic chips used for droplet-based scRNA-seq. In contrast, protoplast-derived plant cells assume a spherical shape, making them more compatible with the microfluidic channels of such platforms. Therefore, for plant tissues and organs with large cells, a well-based scRNA-seq platform such as Rhapsody™ (BD) may provide a more suitable alternative. Additionally, existing bio-sorters such as the Hana Namocell are suitable to cells up to 40 µm in diameter, such as maize meiocytes,^28^ which should accommodate most plant cell types.

Third, FX-Cell, FXcryo-Cell, and cryoFX-Cell do not guarantee successful single-cell preparation for all plant tissues and organs. This limitation likely arises from the presence of specialized cell wall components in certain plants. Further optimization of digestion enzyme combinations and formulations may enhance dissociation efficiency in the future.

Over the past two decades, scRNA-seq has revolutionized the understanding of animal cell identity, development, and evolution, yet its application in plants has been slower to develop due to the extensive optimization required for protoplast isolation. FX-Cell has the potential to overcome these barriers, facilitating discoveries across diverse plant species and tissues. Ultimately, these methods could lay the foundation for the development of Plant Cell Atlases for every target species.^23,24^

## Acknowledgements

We thank Dr. Yiwen Deng (Zhejiang University, China) and Dr. Feng Zhou (CEMPS, CAS) for providing *Oryza longistaminata* plants. This work was supported by the grants from Biological Breeding-National Science and Technology Major Project (2023ZD04073), and National Natural Science Foundation of China (32388201), Strategic Priority Research Program of the Chinese Academy of Sciences (XDB0630201), New Cornerstone Science Foundation through the XPLORER PRIZE, and the National Science Foundation awards (1907220 to D.B.M. and 17540974 to Blake Meyers and V.W.).

## Author contributions

D.B.M. and B.N. initiated the idea of FX-Cell method and generated data on maize anthers and diverse plants samples. X.M. and J.-W.W. improved FX-Cell and then developed FXcryo-Cell and cryoFX-Cell methods. B.N., Y.-Q.W., and X.M. designed and optimized the enzyme RNase-depletion protocol. X.M., H.-C.X., K.L., and Q.-L.S. generated cell atlases. M.-C.W., Z.-G.X., and X.M. performed bioinformatic analyses. Z.-D.Z. and Y.X.H. contributes to rice samples. J.G. prepared figures. D.B.M. and J.-W.W. wrote the manuscript with input of V.W. and X.M..

## Declaration of interests

A patent related to the enzyme RNase-depletion method has been awarded to Stanford University with B.N. as inventor (U.S. Patent No. 11,519,831).

## Extended data tables

**Extended Data Table 1. Statistics results of scRNA-seq and cell type annotation.**

Sheet 1. Statistical summary of scRNA-seq data.

Sheet 2. Marker genes and source for maize anther cell atlas.

Sheet 3. Marker genes and source for rice root cell atlas.

Sheet 4. Marker genes and source for rice tiller node cell atlas.

Sheet 5. Marker genes and source for wild rice rhizome node cell atlas.

Sheet 6. Marker genes and source for maize root crown cell atlas.

Sheet 7. Marker genes and source for Arabidopsis leaf cell atlas.

**Extended Data Table 2. Supporting data for wound response analysis at single-cell resolution.**

Sheet 1. Wound response genes used for AUCell analysis.

Sheet 2. Wound response gene list in bundle sheath cells.

Sheet 3. Wound response gene list in epidermis.

Sheet 4. Wound response gene list in hydathodes.

Sheet 5. Wound response gene list in mesophyll cells.

Sheet 6. Wound response gene list in phloem.

Sheet 7. GO term analysis of unique wound-induced genes in each cell type.

Sheet 8. GO term analysis of unique wound-repressed genes in each cell type.

**Extended Data Table 3. Supporting data for stratified sampling of the cells in the atlas.**

Sheet 1. Stratified cells from published scRNA-seq data of rice root tip.

Sheet 2. Stratified cells from published scRNA-seq data of Arabidopsis leaf.

## Extended data figures and figure legends

**Extended Data Fig. 1.**
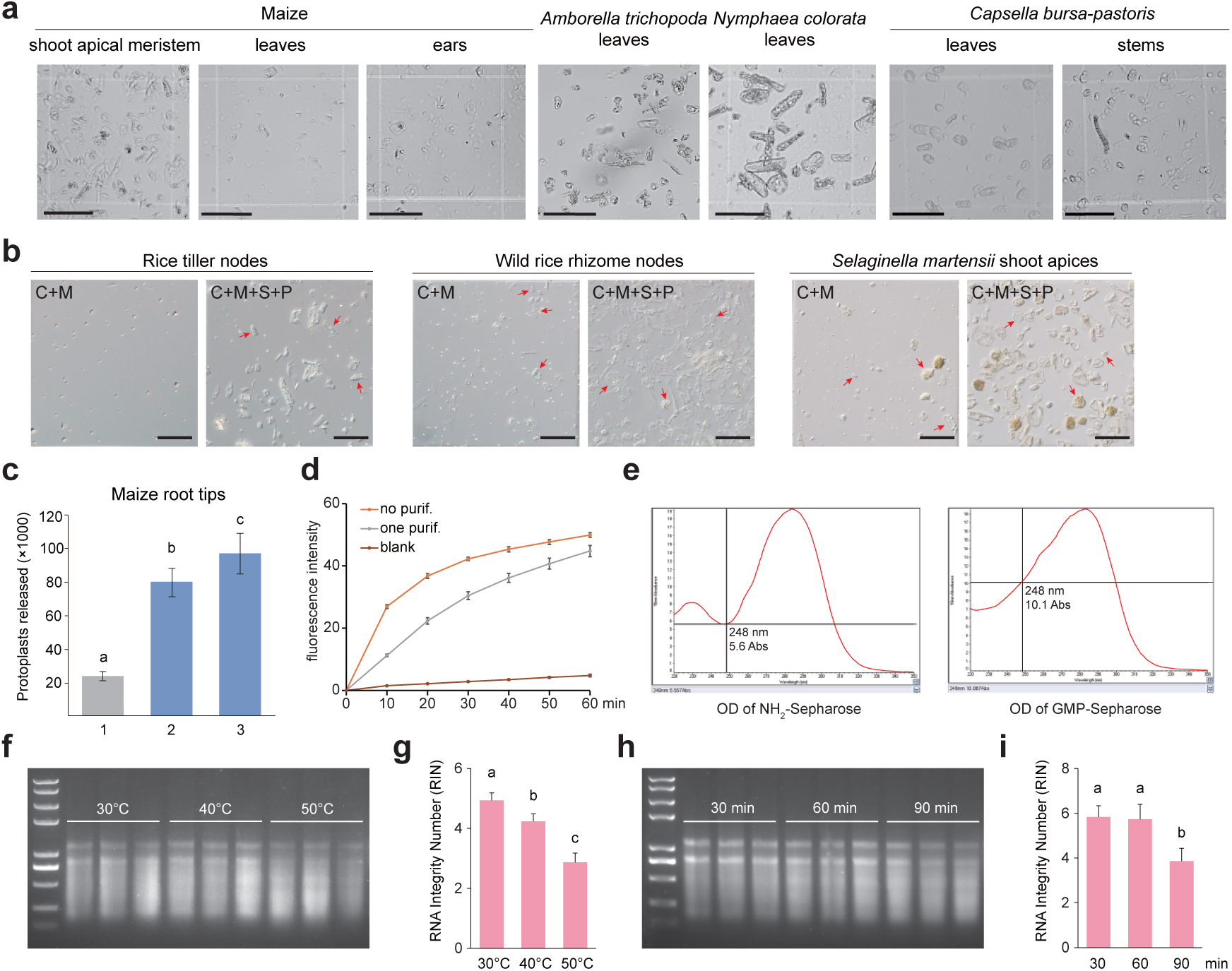
Supporting data for the generation of the fixation and digestion at high temperatures protocol. **a,** Representative image showing the released cells from different tissues and species. Varying plant tissues were digested with the fixation and digestion at high temperatures protocol. See quantification results in Fig. 1c. **b,** Images showing the cells prepared from fixed rice tiller nodes, wild rice rhizome nodes, and shoot apices of *Selaginella martensii*. The tissues were subjected to fixation and digestion at high temperatures using a combination of two enzymes (i.e., reduced enzyme mix, C+M) or four enzymes (C+M+S+P). Red arrows indicate single cells. Please note that the number of released cells was greatly increased when tissues were digested by a combination of four enzymes. Scale bars, 50 μm. See quantification results in Fig. 1d. **c,** Quantification of released protoplasts or cells from maize root tips. We compared the optimized protoplasting protocol from Ortiz-Ramirez et al. (2018)^45^ and our fixation-based protocol. Maize root tips were digested following Ortiz-Ramírez et al. (2018) and released protoplasts were quantified via hemocytometer (grey bar, 1). Maize root tips were fixed then digested at 50°C for 90 minutes using the Ortiz-Ramírez et al. (2018) protoplast enzyme mix (blue bar, 2) or reduced enzyme mix (blue bar, 3). The released cells were similarly quantified. Data are the mean ± s.e.m. from five independent biological replicates. Different letters denote statistically significant variation (Student’s *t*-test, *p* < 0.05). **d,** Assessment of RNase enzymatic activity in the digestion solution. “purif.” indicates that RNase was removed from the digestion solution via affinity chromatography. “Blank” utilizes the enzyme-dissolving RNase binding buffer (RBB) as a control. Data are the mean ± s.e.m. from three independent experiments. **e,** Evaluation of the coupling efficiency of GMP onto NH_2_-Sepharose. The uncoupled NH_2_-Sepharose exhibited an absorbance of 5.6 Abs at 248 nm, while the NH_2_-Sepharose successfully coupled with GMP (GMP-Sepharose) displayed an absorbance of 10.1 Abs at 248 nm. **f,h,** Assessment of RNA integrity through electrophoresis analysis. Rice roots were fixed at various temperatures (**f**) and times (**h**). RNAs were extracted and subjected to electrophoresis analysis. Please note that RNA integrity decreased with increased time and temperature. On the far left is the DNA ladder. **g,i,** Assessment of RNA integrity through RIN analysis. Rice roots were fixed at various temperatures (**g**) and times (**i**). RNAs were extracted and subjected to RIN analysis. Please note that RNA integrity decreased with increased time and temperature. Data are the mean ± s.e.m. from three independent experiments.

**Extended Data Fig. 2.**
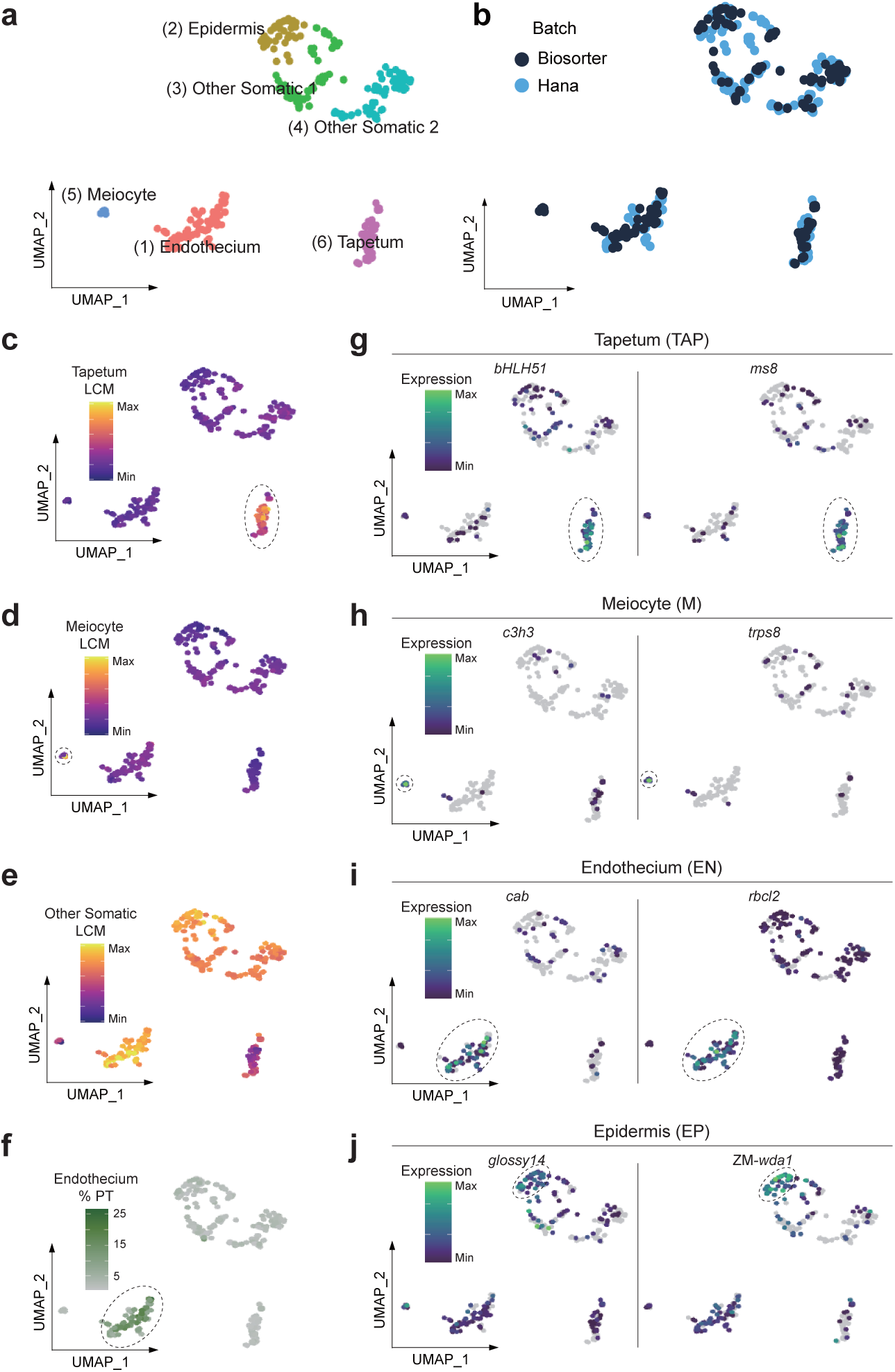
Supporting data for the generation of the maize anther cell atlas by the fixation and digestion at high temperatures protocol and CEL-seq2. **a,** UMAP clustering of 307 cells from 2.0 mm maize anthers. Six distinct clusters are shown in different colors. **b,** Comparison of different cell isolation method (Biosorter vs. Hana). Each dot represents a single cell. **c-e,** Correlation values of each cell with LCM tapetal (**c**), meiocyte (**d**), and other somatic cell types (middle layer, endothecium, and epidermis) (**e**) data. **f,** Percentage of total UMIs originating from the plastid for each cell. Please note that Murphy et al. uncovered that the endothecium contains chloroplasts unlike the other anther cell layers.^55^ **g-j,** UMAP plots showing expression patterns of tapetal (**g**), meiocyte (**h**), putative endothecium (**i**), and putative epidermis (**j**) marker genes. The corresponding cell clusters are labeled by dashed circles.

**Extended Data Fig. 3.**
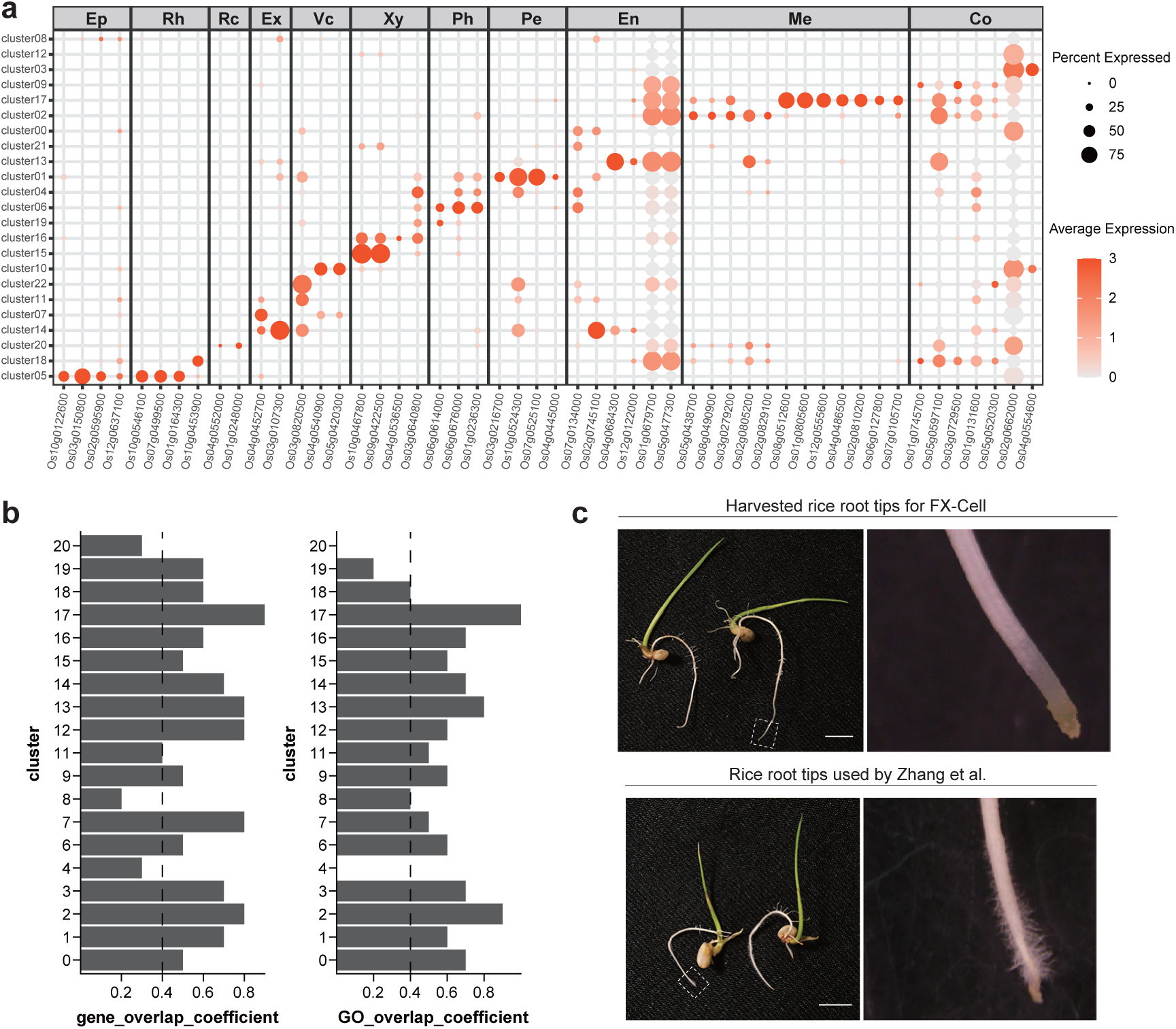
Supporting data for the annotation of the integrated rice root tip cell atlas. **a,** Expression patterns of published cell type marker genes in the integrated rice root tip cell atlas generated by FX-Cell, FXcryo-Cell, and cryoFX-Cell (Fig. 3g; Extended Data Table 1). The diameter of the points represents the proportion of cells expressing a specific gene within a given cluster, while the color of the points indicates the relative expression level of the gene. Ep, epidermis; Rh, root hairs; Rc, root cap; Ex, exodermis; Vc, vascular cylinder; Xy, xylem; Ph, phloem; Pe, pericycle; En, endodermis; Me, meristem-like cells; Co, cortex. **b,** Cluster similarity analysis. The rice root tip cell clusters generated by traditional scRNA-seq and FX-Cell were compared. The overlap ratios of marker genes (left) and GO signaling pathways (right) of each cluster are given. An overlap coefficient greater than or equal to 0.4 (dashed line) is considered indicative of high similarity. **c,** Representative images showing the harvested rice root tips (white dashed boxes) for FX-Cell (Fig. 2e). The rice seeds were geminated and cultured on plates. Please note that, compared to the sample used by Zhang et al.,^34^ the sample used for FX-Cell developed fewer root hairs (magnified images on the right side) due to the increased water availability in the plates. Scale bars, 0.5 cm.

**Extended Data Fig. 4.**
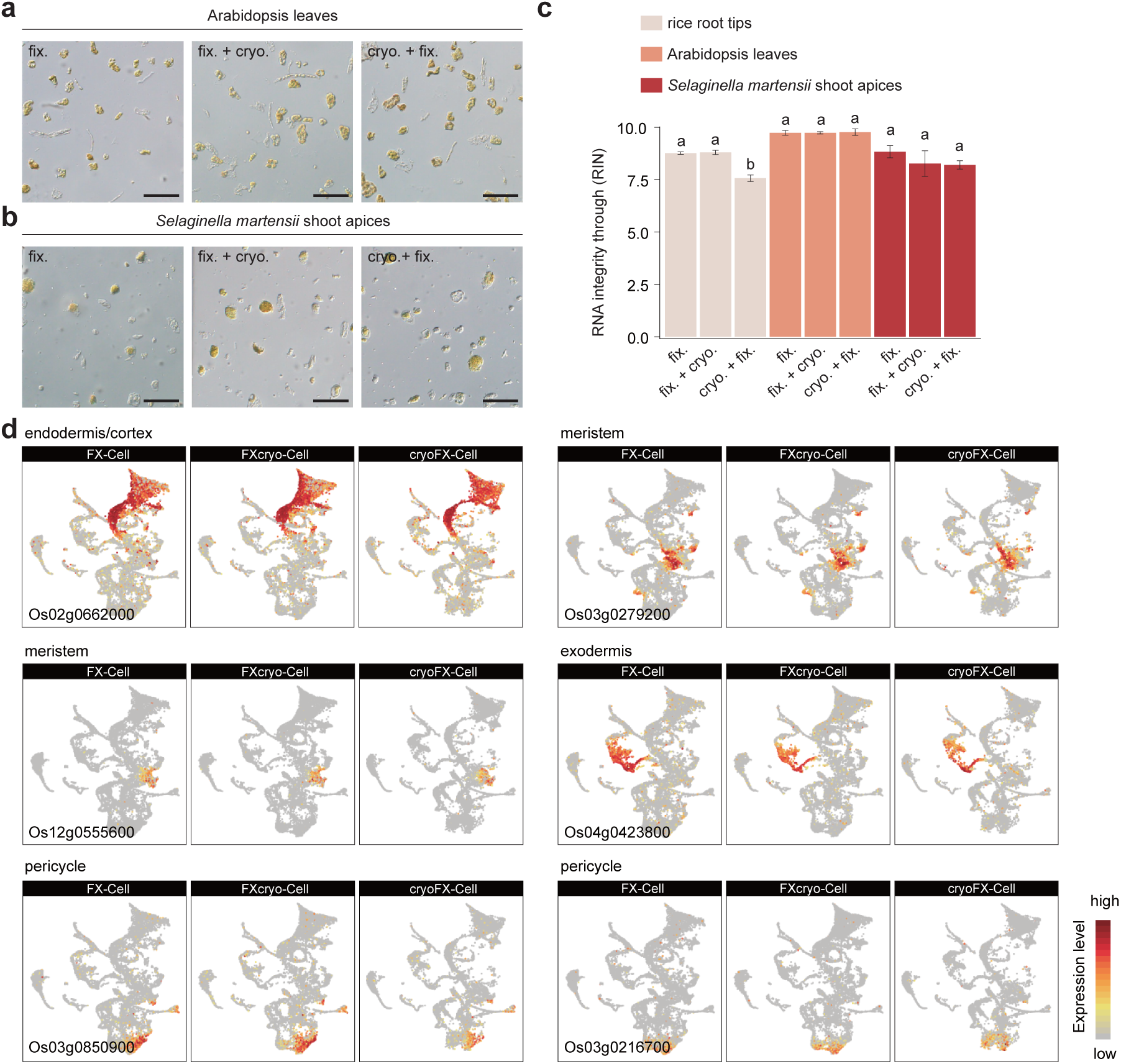
Supporting data for the comparison of rice root tip cell atlases generated by different scRNA-seq methods. **a,b,** Images showing the cells derived from the leaves of *A. thaliana* (**a**) and shoot apices of *S. martensii* (**b**) with three different methods. “fix.” indicates the samples that were fixed and subsequently subjected to high-temperature enzymatic digestion; “fix. + cryo.” refers to samples that were fixed, frozen, cryopreserved, then thawed and subjected to high-temperature enzymatic digestion; “cryo. + fix.” indicates the samples that were directly frozen and cryopreserved, then thawed, fixed, and subjected to high-temperature enzymatic digestion. Scale bars, 50 μm. **c,** Assessment of RNA integrity through RIN analysis. Samples were fixed and cryopreserved as described in (**a,b**). Data are the mean ± s.e.m. from three independent biological replicates. Different letters denote statistically significant variation (Student’s *t*-test, *p* < 0.05). **d,** UMAP plots showing the expression patterns of other cell type marker genes of rice root tip cell atlases generated by FX-Cell, FXcryo-Cell, or cryoFX-Cell (Fig. 2e; Figs. 3d,e).

**Extended Data Fig. 5.**
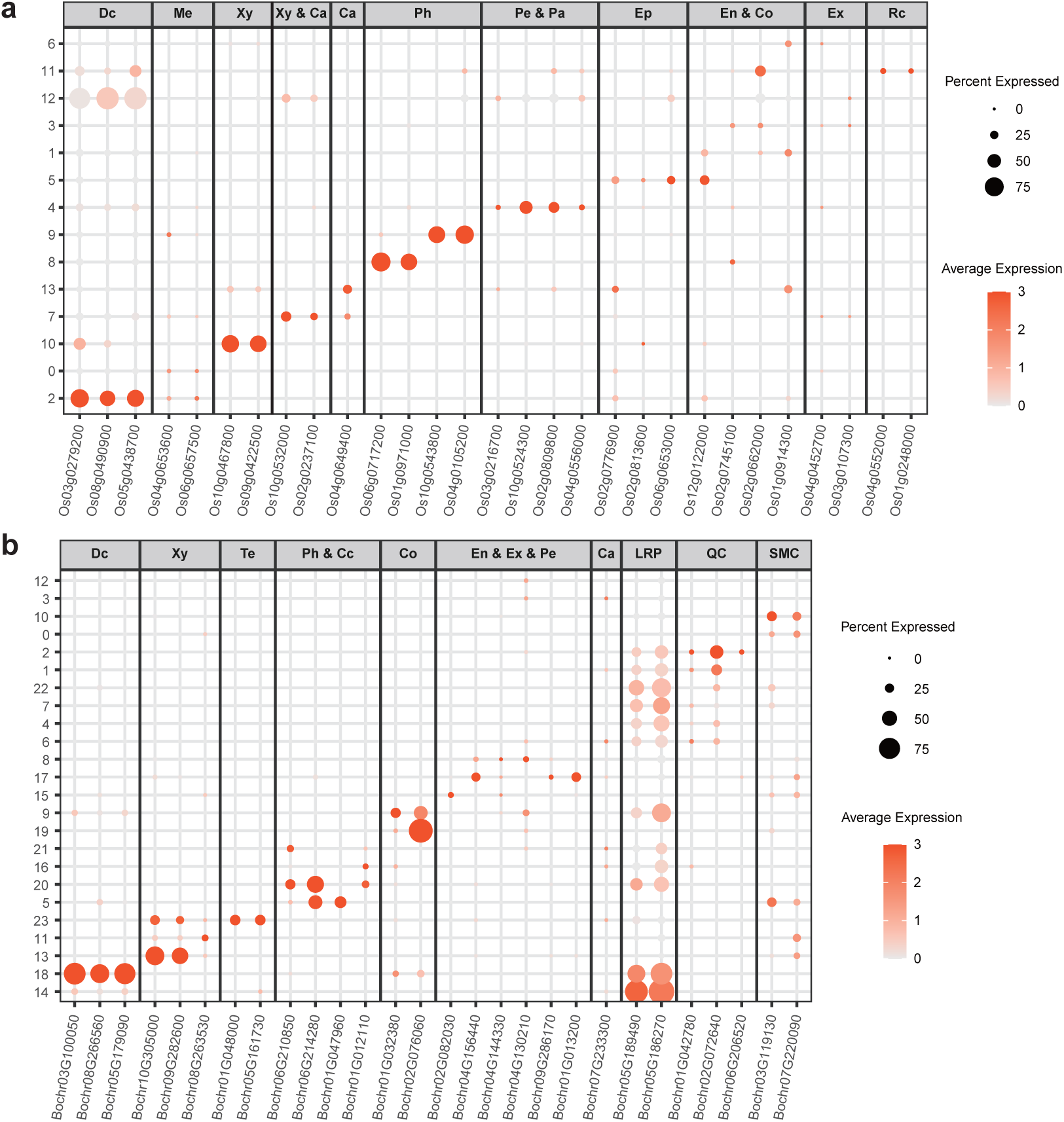
Supporting data for the cell atlases of difficult-to-digest rice samples. **a,** Expression patterns of cell type marker genes in the rice tiller node cell atlas generated by FXcryo-Cell (Fig. 4a; Extended Data Table 1). The diameter of the dots represents the proportion of cells expressing a specific gene within a given cluster, while the color of the dots indicates the relative expression level of the gene. Dc, dividing cells; Me, meristem-like; Xy, xylem; Xy & Ca, xylem/cambium; Ca, cambium; Ph, phloem; Pe & Pa, pericycle & parenchyma; Ep, epidermis; En & Co, endodermis & cortex; Ex, exodermis; Rc, root cap. **b,** Expression patterns of cell type marker genes in the wild rice rhizome node cell atlas generated by FXcryo-Cell (Fig. 4b; Extended Data Table 1). The diameter of the dots represents the proportion of cells expressing a specific gene within a given cluster, while the color of the dots indicates the relative expression level of the gene. Dc, dividing cells; Xy, xylem; Te, tracheary elements; Ph & Cc, phloem & companion cells; Co, cortex; En & Ex & Pe, endodermis, exodermis, and pericycle; Ca, cambium; LRP, lateral root primordia; QC, root quiescent center; SMC, shoot meristem like.

**Extended Data Fig. 6.**
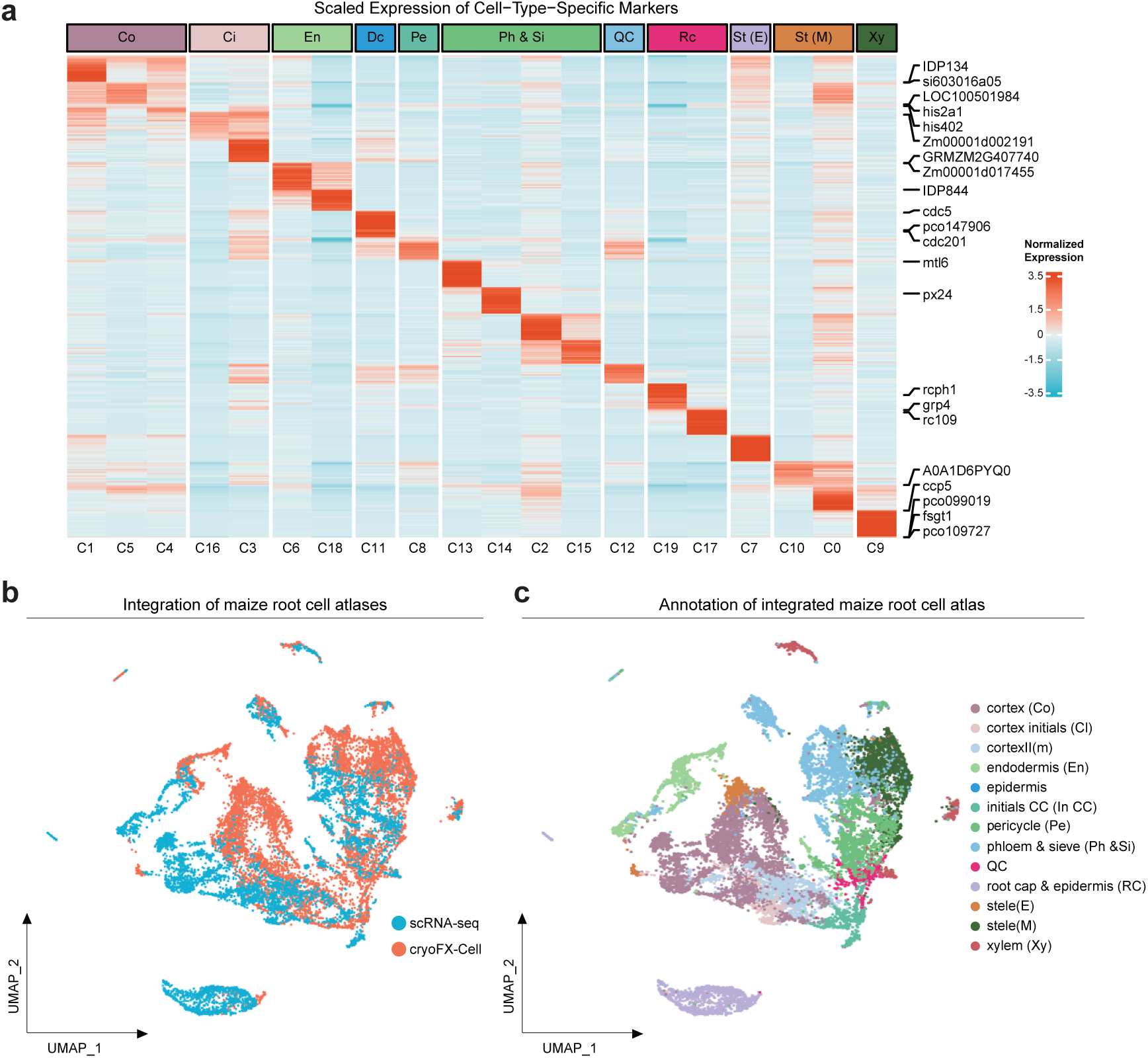
Supporting data for the cell atlases of field-grown maize sample. **a,** Heatmap showing the top 150 genes from each cluster in the maize crown root cell atlas generated by cryoFX-Cell (Fig. 4d). Cell types are annotated using published maize root marker genes (Extended Data Table 1). The representative genes in each cell type are given on the right. Normalized expression levels are shown. Co, cortex; Ci, cortex initials; En, endodermis; Dc, dividing cells; Pe, pericycle; Ph & Si, phloem & sieve; QC, root quiescent center; Rc, root cap; St, stele (E); St, stele (M); Xy, xylem. **b,** Joint analysis of the maize root cell atlas generated by cryoFX-Cell (Fig. 4d) and traditional scRNA-seq.^45^ Harmony algorithm was applied to mitigate batch effects. **c,** Annotation of the integrated maize root tip cell atlas shown in (**b**), based on the published cell type marker gene (Extended Data Table 1). Different colors represent different cell types.

**Extended Data Fig. 7.**
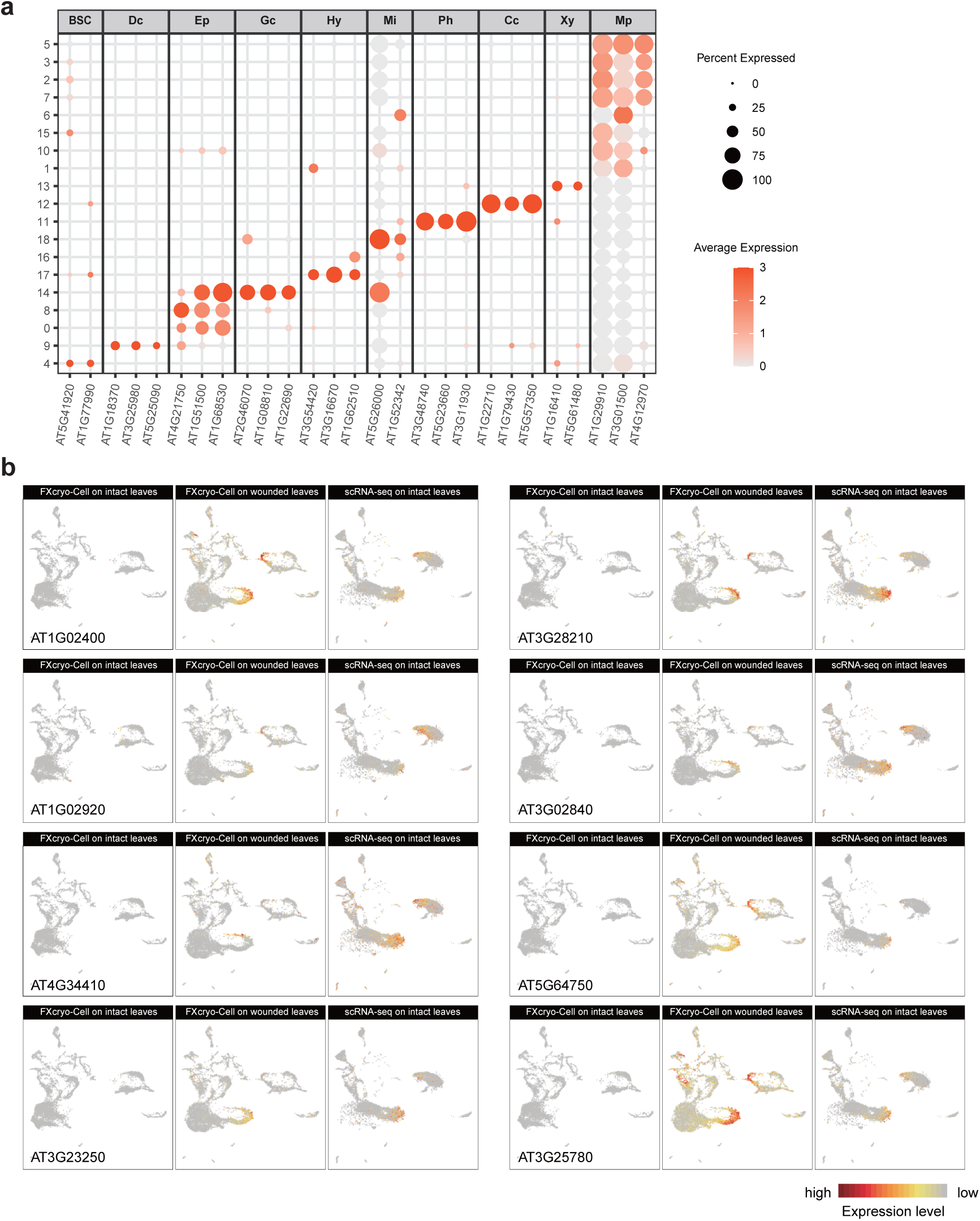
Supporting data for annotation of *A. thaliana* leaf cell atlas and analysis of wound response genes. **a,** Expression patterns of representative cell type marker genes in each cell cluster of the integrated Arabidopsis leaf cell atlas in Fig. 5b (Extended Data Table 1). The diameter of the dots indicates the proportion of cells expressing a specific gene within a given cluster, while the color of the dots represents the relative expression levels of the genes. BSC, bundle sheath cells; Dc, dividing cells; Ep, epidermis; Gc, guard cells; Hy: hydathodes; Mi, myrosin idioblasts; Ph, phloem; Cc, companion cells; Xy, xylem; Mp, mesophyll. **b,** UMAP plots showing the expression patterns of representative wound-induced genes^52^ in the integrated cell atlas in Fig. 5b.

**Extended Data Fig. 8.**
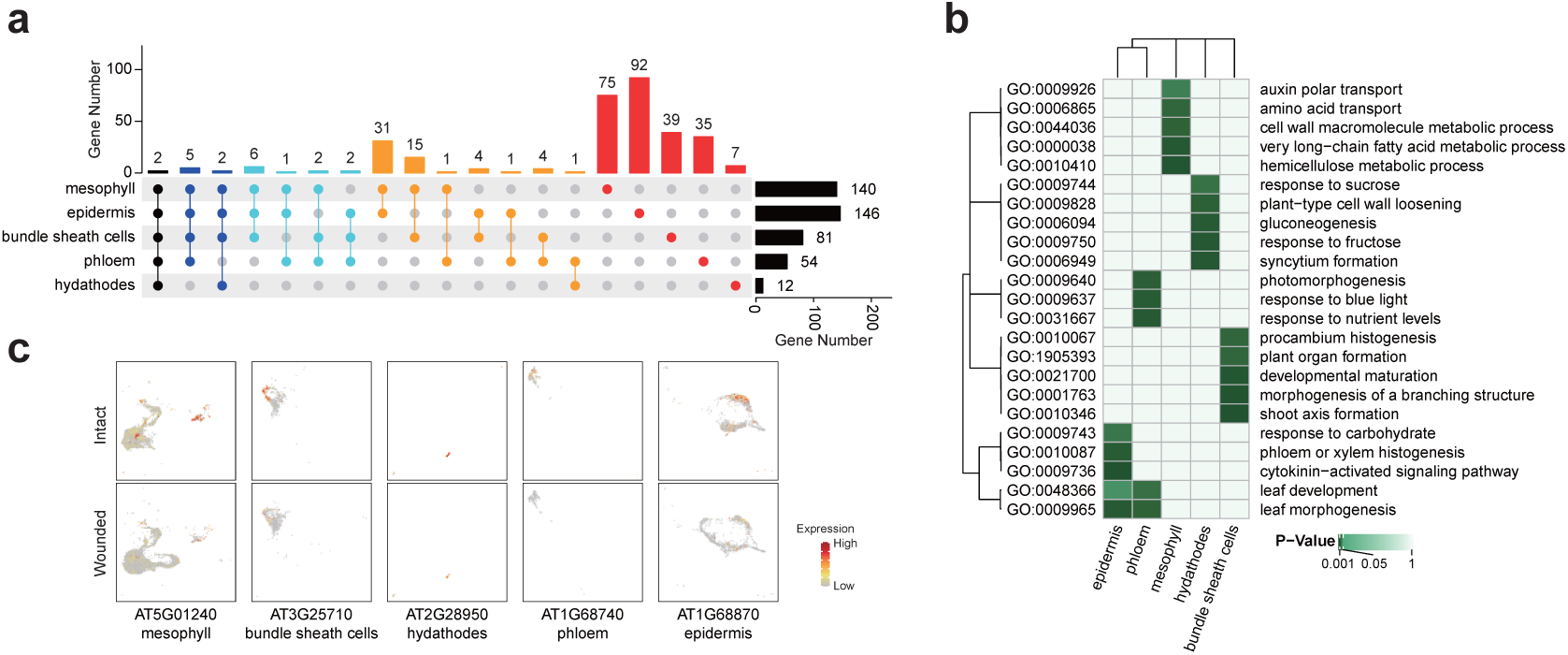
Supporting data for cell-type specific wound-response genes. **a,** Upset plot presenting shared and unique wound-repressed genes in each cell type. The number of genes for each cell type is presented on the top of the colored bars. Total number of wound-induced genes in each cell type is presented on the right. **b,** GO term analysis of unique wound-repressed genes in each cell type. **c,** UMAP plots presenting representative cell-type specific wound-repressed genes. Only cells corresponding to each specific cell type are presented. The upper and lower rows indicate intact and wounded *A. thaliana* leaf cell atlases produced using FXcryo-Cell (Fig. 5c), respectively.

**Extended Data Fig. 9.**
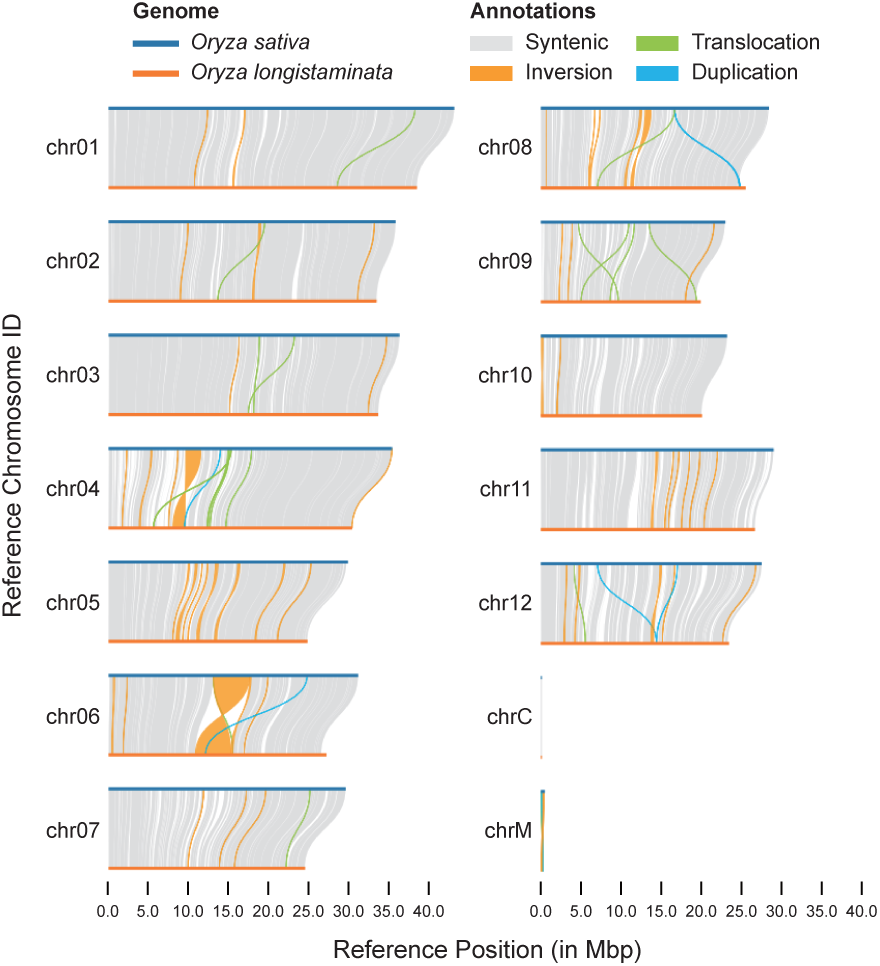
Supporting data for the genome assembly of *O. longistaminata*. The genomic differences between two genome assemblies, *Oryza sativa* and *Oryza longistaminata*, are illustrated by plot using plotsv.

## Methods

### Plant materials and growth conditions

For maize anther, root tip, shoot apical meristem, leaf, and ear samples (Figs. 1a,b,c; Extended Data Figs. 1a,c), *Zea mays* (inbred line W23 bz2) individuals were grown under greenhouse conditions in Stanford, CA, USA (14 hours light/10 hours dark). Daily irrigation and fertilization were maintained for robust growth. Beginning five to six weeks after planting, individual plants were felled ∼20 cm above ground level for anther dissection between 8:00 and 9:00 am. The sacrificed plants were taken to the lab within 10 minutes where the tassels were dissected out of the stem and leaf whorl. For maize crown root sample (Fig. 4d), *Z. mays* (inbred line B73) were germinated in the greenhouse, and the 2-week-old seedlings were transplanted to the field in Shanghai. The crown roots were harvested from 45-day-old plants.

The seeds of *Oryza sativa* (ZH11) were sown on filter paper soaked in water in petri dishes. After 5 days at 28°C (10 hours light/14 hours dark), the root tips (Extended Data Fig. 3c) and tiller nodes (insets in Fig. 4a) were dissected and digested. *Oryza longistaminata* (wild rice) seedlings were grown in a greenhouse at 28°C (12 hours light/12 hours dark). The rhizome nodes were harvested for digestion (insets in Fig. 4b). *Selaginella martensii* seedlings were purchased from the market and cultivated in the greenhouse at 21°C (16 hours light/8 hours dark). The shoot apices were harvested for digestion. The seeds of *Arabidopsis thaliana* (Col-0) plants were grown in the greenhouse at 21°C (16 hours light/8 hours dark), and the third-to-last true leaves of 24-day-old plants were harvested for digestion (intact sample). For wounding treatment, the leaves were wounded by a syringe needle and harvested for digestion (wounded sample). *Amborella trichopoda* and *Nymphaea colorata* were grown under tropical (27°C) greenhouse conditions in Stanford, CA, USA (12 hours light/12 hours dark).

### Maize anther dissection and protoplasts/cells preparation

A Leica M60 dissecting scope and stage micrometer (ThermoFisher Scientific) were used to isolate 2.0 mm anthers from the upper florets of spikelets along the central spike of the tassel. Protoplasts/cells release of fixed and fresh maize anthers was compared in a variety of conditions. Three 2.0 mm anthers were pooled per replicate with five replicates per condition. Fresh anthers were digested at 30°C for 90 minutes or 16 hours in the enzyme mix (1.25% w/v Cellulase-RS, 0.5% Pectolyase Y-23, 0.5% Macerozyme-R10, 0.5% Hemicellulose) from Nelms and Walbot.^28^ Fixed samples were left in ice-cold Farmer’s solution (3:1 100% ethanol:glacial acetic acid) for 2 hours, washed twice in ice-cold 0.1× phosphate-buffered saline (PBS; Sigma-Aldrich) for 5 minutes, then digested at 30°C or 50°C for 90 minutes in 20 mM MES, pH 5.7 with a 1:10 dilution of RNase-depleted reduced enzyme mix stock (enzymes were stored in glycerol stocks, see below; stocks were normalized so that a 1:10 dilution has the same A280 as a 1.25% w/v Cellulase-RS and 0.4% w/v Macerozyme-R10 solution). The protoplasts/cells from the digested, fixed anthers were dissociated via shear force between two microscope slides with thin tape as a spacer. For each replicate, the number of protoplasts/cells was estimated using a hemocytometer and then averaged. Images of the dissociated cells were taken on a Nikon Diaphot inverted microscope with a mounted Nikon D40 camera.

### Protoplasts/cells preparation from other tissues and species

Protoplasts/cells release from maize root tips was compared in three conditions (Extended Data Fig. 1c): 1) fresh protoplasting as described,^45^ 2) fixation and digestion with the enzyme concentrations as described,^45^ or 3) fixation and digestion with reduced enzyme mix. Maize seeds were treated, germinated, and grown as described.^45^ Seedling primary roots were cut 5 mm above the tip with a scalpel. Three root tips were pooled per replicate with five replicates per condition. Fresh root tips were pre-treated, washed, digested in the enzyme mix as described,^45^ filtered, and washed. The fresh protoplasts were counted with a hemocytometer. The fixed samples were left in ice-cold Farmer’s solution for 2 hours, washed twice with ice-cold 0.1× PBS for 5 minutes then digested at 50°C with the enzyme mix as described^45^ or reduced enzyme mix as described above (Extended Data Fig. 1c). Digested tissue was manually disrupted with pipetting, then the number of individual cells counted with a hemocytometer.

We expanded our sampling by comparing our method to a standard protoplasting protocol^28^ in three additional maize tissues (apical meristems, young leaves, young ears) and four non-model plant taxa and tissues (leaves from the basal angiosperms, *Amborella trichopoda* and waterlily, *Nymphaea colorata*, leaf and stem tissue from the non-model Brassicaceae *Capsella bursa-pastoris*) (Fig. 1c). For each tissue, comparably sized samples were either fixed in Farmer’s solution, washed twice 0.1× PBS, and digested at 50°C with reduced enzyme mix or directly digested at 30°C for 90 minutes in the enzyme mix from Nelms and Walbot.^28^ For each replicate, the number of protoplasts/cells was estimated using a hemocytometer then averaged. Images of the dissociated cells were taken on a Nikon Diaphot inverted microscope with a mounted Nikon D40 camera.

### Single cell preparation from rice tiller nodes, wild rice rhizome nodes, and shoot apices of *S. martensii*

Plant samples were harvested in ice-cold Farmer’s solution and fixed for 2 hours, followed by two washes in ice-cold 0.1× PBS for 5 minutes each. The plant samples were then shredded and digested at 50°C for 90 minutes in either C+M cocktail solution (i.e., reduced enzyme mix, 1.25% Cellulase-RS, 0.4% Macerozyme-R10, 0.4 M mannitol, 20 mM 4-morpholineethanesulfonic acid, pH 5.7, 10 mM KCl, 10 mM CaCl_2_, and 0.1% BSA) or C+M+S+P cocktail solution (1% Cellulase-RS, 0.5% Macerozyme-R10, 1% Snailase, and 0.5% Pectinase, 0.4 M mannitol, 20 mM 4-morpholineethanesulfonic acid, pH 5.7, 10 mM KCl, 10 mM CaCl_2_, and 0.1% BSA). The samples were gently sucked and blown to fully release the cells. The cells were filtered twice using cell strainers (40 μm diameter, Falcon, Cat No./ID: 352340), spun at 300g for 3 minutes, and washed three times with 0.1× PBS at room temperature (RT). The concentration of cells was counted using a hemocytometer (Fig. 1d; Extended Data Fig. 1b).

### Preparation of GMP-Sepharose/Agarose

To prepare the GMP-Sepharose, we proceeded as follows: Add 1.028 g NaIO_4_ to 24 mL of ultrapure H_2_O (i.e., 0.2 M NaIO_4_). After NaIO_4_ is completely dissolved, add 1.952 g guanosine 5′-monophosphate disodium salt (5’-GMP), and incubate at RT, protected from light, with gentle shaking for 1 hour. The resultant solution is called oxidized GMP solution.

Resuspend 25 mL NH_2_-Sepharose (Shanghai Puke, Cat No./ID: RF1052) by 50 mL of ultrapure H_2_O, and then filter to remove the liquid with the Büchner funnel. Repeat this step twice more. Add 50 mL of 0.1 M borax (pH 9.0), thoroughly suspend NH_2_-Sepharose, and filter to remove the liquid with the Büchner funnel. Repeat this step two more times. Add 72 mL of 0.1 M borax (pH 9.0), fully resuspend NH_2_-Sepharose, and transfer it to a glass beaker. Wash the Büchner funnel with another 72 mL of 0.1 M borax (pH 9.0) once, and transfer the liquid to the same beaker. Take 3 mL of NH_2_-Sepharose and add 0.5 mL of 0.2 M GMP (non-oxidized GMP). Mix thoroughly by shaking and place it at 4°C (this sample serves as a negative control for evaluation of the coupling efficiency).

Add 200 μL of 50% glycerol to the aforementioned oxidized GMP solution and incubate at RT with gentle shaking for 30 minutes. Add 24 mL of this solution to 141 mL of NH_2_-Sepharose, mix thoroughly to suspend NH_2_-Sepharose, cover the beaker with plastic wrap, and incubate at RT with gentle shaking for 2 to 4 hours. At this point, NH_2_-Sepharose has initially coupled with GMP, and we refer to it as GMP-Sepharose thereafter.

Transfer the GMP-Sepharose to a Büchner funnel to remove the liquid. Add 50 mL of 0.1 M borax (pH 9.0), fully suspend GMP-Sepharose, and filter to remove the liquid. Repeat this step two more times. Add 84 mL of 0.1 M borax (pH 9.0), thoroughly shake to resuspend GMP-Sepharose, and transfer it to a glass beaker. Wash the Büchner funnel with another 84 mL of 0.1 M borax (pH 9.0) once, and transfer the liquid to the same beaker.

Slowly add 544 mg NaBH_4_, incubate at 4°C with gentle shaking for 1 hour (cover with plastic wrap, poking a small hole to ensure ventilation), and remove the liquid with the Büchner funnel. Wash the GMP-Sepharose three times with 50 mL of 0.1 M borax (pH 9.0), once with 50 mL of H_2_O, once with 1 M NaCl, and finally remove the liquid with the Büchner funnel. Transfer GMP-Sepharose out and dilute it to 50 mL with 1 M NaCl for long-term storage. The same procedure can be used to prepare the GMP-Agarose.

### Evaluation of the coupling efficiency of GMP-Sepharose

To evaluate the coupling efficiency of GMP-Sepharose, we proceeded as follows: Take 1 mL of GMP-Sepharose, remove the liquid by centrifuge, and do the same with the NH_2_-Sepharose incubated with non-oxidized GMP. Transfer both to separate 50 mL centrifuge tubes, add 40 mL of H_2_O, and incubate at 4°C with rotation for 30 minutes, then centrifuge to collect the supernatant. Wash GMP-Sepharose and NH_2_-Sepharose with H_2_O twice, using 40 mL each time. Take approximately 100 μL of solid GMP-Sepharose and NH_2_-Sepharose, add 0.5 mL of 1 M HCl, and heat in a boiling water bath for 1 hour. Once the samples have cooled, use a Nanodrop 2000 to measure the DNA concentration, detecting the absorbance in the mode for DNA (using 1 M HCl as the blank). The high-temperature acid hydrolysis of GMP-Sepharose will produce guanine, which has the maximum absorbance at 248 nm. The GMP-coupled NH_2_-Sepharose will show significantly higher absorbance at 248 nm compared to the control NH_2_-Sepharose (Extended Data Fig. 1e).

### Depletion RNase from digestion enzymes using GMP-Sepharose/Agarose

To deplete RNase from digestion enzymes, we proceeded as follows: Resuspend the cell digestion enzymes in RNase binding buffer (RBB; 150 mM NaCl, 10 mM citrate, pH 7.0). For the samples in Figs. 1b,c,f, we used C+M cocktail. For other samples, we used the C+M+S+P cocktail. Spun at 5,000 rpm at 4℃ for 5 minutes, collect the supernatant. Repeat this step once. Take 100 μL of these enzyme solution for later use and measure the protein concentration using the Protein A280 mode on a Nanodrop 2000.

We used a two-step method involving gravity chromatography column and FPLC column to purify the digestion enzymes: Load 4 mL of GMP-Sepharose onto a gravity chromatography column. Fill the remaining volume with 21 mL of GMP-Sepharose in an FPLC column (Aogma, Cat No./ID: XK-1030A). Equilibrate the GMP-Sepharose gravity chromatography column three times with 5 mL of RBB. Slowly add the enzyme solution to the GMP-Sepharose gravity chromatography column, collecting the effluent. After all the enzyme solution has passed through the resin, slowly add an additional 6 mL of RBB to the resin surface and collect the effluent. Transfer the effluent to a Sartorius 15 mL 10 kDa MWCO PES ultrafiltration tube (Sartorius, Cat No./ID: FUF151) and centrifuge at 4,000g, 4°C to concentrate the protein. When the protein concentration approaches that of the original enzyme solution, transfer it to a 2 mL centrifuge tube. To regenerate the column, wash the GMP-Sepharose gravity chromatography column three times with 5 mL of ultrapure H_2_O, three times with 5 mL of RNAse elution buffer (5 M NaCl, 1 mM 5’-GMP, 1 mM 2’(3’)-GMP), three times with 5 mL of 1 M NaCl. Add 4 mL of 1 M NaCl and store the GMP-Sepharose column at 4°C for repeated use.

Equilibrate the GMP-Sepharose FPLC column with 45 mL of RBB. Load 1 mL the concentrated enzyme onto the GMP-Sepharose FPLC column. Collect fractions and pool those with A280 absorbance > 0.1. Concentrate pooled enzymes with Sartorius 15 mL 10 kDa MWCO PES ultrafiltration tube. When the protein concentration reaches that of the original enzyme solution, transfer it to a 2 mL centrifuge tube. To regenerate the GMP-Sepharose FPLC column, wash the column with 45 mL RNAse elution buffer and 45 mL 1 M NaCl. To further reduce RNAse (Fig. 2d), the obtained enzyme solution can be purified again using a GMP-Sepharose FPLC column. Finally, we obtain a 10× RNase-depleted digestion enzyme stock solution, at the same concentration as the original solution.

### RNase activity assays

To prepare the enzyme solution, the 10× original enzyme or 10× RNase-depleted enzyme was diluted to a 1× concentration using digestion buffer (0.4 M mannitol, 20 mM MES, pH 5.7, 10 mM KCl, 10 mM CaCl_2_, and 0.1% BSA). RNase inhibitor (1 μL/μL, ThermoFisher Scientific, Cat No./ID: N8080119), tRNA (1 μg/μL), and triGMP (1 mM) were then added, as illustrated in Fig. 2d. RBB buffer was used as a control. RNase activity was quantified using the RNAse Activity Fluorescence Detection Kit (Biyuntian, Cat No./ID: P0347S) (Fig. 2d; Extended Data Fig. 1d).

### Assessment of RNA integrity

Maize anther RNA quality was tested in four conditions of cell preparation (Fig. 1f): 1) flash frozen, 2) fixed in Farmer’s solution then washed twice in 0.1× PBS then flash frozen, 3) fixed, washed, and digested in commercial enzymes, and 4) fixed, washed, and digested in RNase-depleted enzymes purified by GMP-Agarose. For each condition, 2.0 mm anthers were isolated from five separate plants with ten anthers pooled per plant. The flash frozen samples were homogenized via bead beating in a 2000 Geno/Grinder (SPEX CertiPrep) with baked 4 mm steel balls. The fixed samples were left in ice-cold Farmer’s solution for 2 hours, washed twice in ice-cold 0.1× PBS for 5 minutes, then incubated at 50°C for 90 minutes with RNase-depleted or commercial enzymes (1.25% w/v Cellulase-RS and 0.4% w/v Macerozyme R10). The RNeasy Plant Mini Kit (Qiagen) was used to extract RNA from samples via the standard protocol. RNA was quality-checked on an Agilent 2100 BioAnalyzer with the RNA 6000 Nano assay (Agilent Technologies). The RNA Integrity Number (RIN) for the five replicates of each condition were averaged and reported alongside the error.

Using a similar protocol, we examined the RNA qualities of the rice root tip samples that were fixed and digested with the C+M cocktail at different temperatures and times (Extended Data Figs. 1g,i), as well as the samples prepared by different methods (Extended Data Fig. 4c). Additionally, we assessed RNA integrity through RNA gel electrophoresis (Extended Data Figs. 1f,h).

### Fixed maize anther cell preparation for CEL-Seq2

Anthers from four individuals of wild-type maize (inbred line W23 bz2) were dissected out. One of the three anthers per floret was used for imaging on a Nikon Diaphot inverted microscope with a Nikon D40 mounted camera at 10× magnification. The remaining two anthers per floret were fixed in ice-cold Farmers solution for 2 hours, washed twice for 5 minutes in 0.1× PBS, and then one anther was digested at 50°C for 90 minutes in the RNase-depleted enzyme mix while the other anther was saved at −20°C. Following digestion, shear force was applied to the anther between two microscope slides with thin tape on each end to prevent the anther from being fully crushed. The top microscope slide was slid back and forth 5-10 times and the sample checked under the dissecting scope to ensure separation of the fixed cells. The cells were washed from the slides into 1 mL of cold 0.1× PBS via pipette and stained with SYBR Green I nucleic acid gel stain (Invitrogen) for 20 minutes. The cells were then filtered through a 100 μm (if bound for the BioSorter) or 40 μm (if bound for the Hana) nylon cell strainer (Corning) into 50 mL Falcon tubes. The stained cells were then sorted into 384-well plates or 96-well plates, each well containing 0.8 μL Primer Master Mix (0.225% Triton X-100, 1.6 mM dNTP mix, 1.875 μM barcoded oligo[dT] CEL-Seq2 primers; Sigma-Aldrich, New England Biolabs) using a BioSorter (Union BioMetrica) or Hana Single Cell Dispenser (Namocell). Following cell sorting, the plates were spun at 400g then stored at −80°C.

### CEL-Seq2 library preparation and data analysis

Single cell libraries were prepared following the CEL-Seq2 protocol with modifications.^33^ Read filtering, mapping, initial processing, and cell clustering for CEL-Seq2 dataset were performed as described.^33^ To initially compare our dataset with that of known cell types, we assessed the similarity of our data with laser-capture microdissection (LCM) sequencing data of known cell types and whole anthers,^56^ which were also prepared from 2.0 mm W23 maize anthers using the same CEL-Seq2 library preparation. UMIs were normalized into transcripts per million (TPM) and log transformed after adding a pseudocount of 100. We then subtracted the single cell TPMs by the log transformed TPMs of the whole anthers to produce ratio measurements. The LCM data had samples for tapetal, meiocyte, and other somatic (middle layer, endothecium, epidermis) cell types and were similarly processed relative to the whole anther data. We then calculated the cell-to-cell Pearson’s correlations of all our single cells relative to each of the LCM samples.

Clusters were determined and visualized with UMAP with a resolution of 0.01 (Extended Data Fig. 2a). The data generated by the cells sorted by BioSorter or Hana were highly similar (Extended Data Fig. 2b). Correlation values of each cell with the LCM tapetal, meiocyte, and other somatic cell types were mapped onto the UMAP (Extended Data Figs. 2c-e), as well as the percentages of transcripts from the plastid genome and mitochondrial genome (Extended Data Fig. 2f). The meiocyte cluster was manually separated from the endothecium cluster based on the LCM correlation data and meiocyte marker genes; it is likely that Monocle did not separate these clusters due to the scarcity of meiocyte cells despite the clear separation in the UMAP. The other somatic 1 (OS1) cluster was subset and reclustered to identify and separate the epidermis cluster based on putative marker genes of the known biology of the cell type (Extended Data Figs. 2g-j).

### snRNA-seq on rice root tips

To generate snRNA-seq data for rice root tips (Fig. 2), we followed these steps: First, prepare a 1× NIB buffer from the 4× NIB buffer stock solution (Sigma-Aldrich, Cat No./ID: CELLYTPN1), and add 1× cocktail (Roche, Cat No./ID: 4693132001) along with 0.2 U/μL RNase inhibitor (ThermoFisher Scientific, Cat No./ID: N8080119). Homogenize rice root tips (1.0 cm in length) using a blade in 200 μL of 1× NIB buffer. Transfer the homogenate to a 15 mL conical tube and adjust the volume to 2 mL by adding 1× NIB buffer. Incubate the mixture at 4°C with rotation for 15 minutes. Next, filter the suspension through a 40 μm cell strainer into a 15 mL conical tube. Add 100 μL of 10% Triton X-100 and mix thoroughly. Centrifuge at 1,000g at 4°C for 10 minutes, carefully discarding the supernatant. Resuspend the pellet in 1 mL of 1× PBS (supplemented with 1× cocktail, 0.1% BSA, and 0.2 U/μL RNase inhibitor) and add DAPI (4’,6-diamidino-2-phenylindole) to achieve a final concentration of 1 μg/mL. Perform fluorescence activated nucleus sorting (FANS), loading 200 μL of 1× PBS (supplemented with 1× cocktail, 0.1% BSA, and 0.2 U/μL RNase inhibitor) as the landing buffer. Sort 200,000 nuclei, then centrifuge at 1,000g at 4°C for 10 minutes. Discard the supernatant, add 30 μL of 1× PBS (supplemented with 1× cocktail, 0.1% BSA, and 0.2 U/μL RNase inhibitor), and resuspend the nuclei. Count the number of nuclei using a microscope, and conduct snRNA-seq using the Chromium Next GEM Single Cell 3ʹ Reagent Kits v3.1 (10× Genomics, Cat No./ID: PN-1000268), following the manufacturer’s instructions. Libraries are sequenced in a 150-bp paired-end mode on a DNBSEQ-T7 sequencer (GenePlus, Beijing, China).

### FX-Cell on rice root tips

To generate cell atlas of rice root tips by FX-Cell (Fig. 2), we proceeded as follows: Place the rice root tips (1 cm in length) into Farmers solution in a 2 mL centrifuge tube and apply vacuum until the material settles to the bottom of the tube. Incubate on ice for 30 minutes. Add pre-chilled 0.1× PBS to the fixed material, mix well, and incubate on ice for 5 minutes. Discard the supernatant. Repeat this step once. Prepare the cell wall digestion enzyme solution with the following components: 0.4 M mannitol, 20 mM MES (pH 5.7), 10 mM KCl, 10 mM CaCl_2_, 0.1% BSA, 1× RNase-depleted C+M+S+P digestion enzymes (diluted from aforementioned 10× RNase-depleted digestion enzyme stock solution), 1 mg/mL tRNA, and 1 mM tri-GMP (a mixture of 0.1 M 5’-GMP and 0.1 M 2’(3’)-GMP). Filter the enzyme solution using a 0.45 μm filter. Transfer the material to a small Petri dish (diameter 30 mm), add an appropriate amount of enzyme solution, and incubate with shaking at 40°C and 80 rpm for 10 minutes. Disrupt the material and incubate with shaking at 40°C and 80 rpm for 20 minutes. Filter the material through a 40 μm cell strainer into a 2 mL centrifuge tube, centrifuge at 4°C and 300g for 3 minutes, and discard the supernatant. Resuspend the cells in 1 mL of 0.1× PBS, filter the cell suspension through a 40 μm cell strainer into a 2 mL centrifuge tube, centrifuge at 4°C and 300 g for 3 minutes, and discard the supernatant. Wash the cells with 1 mL of 0.1× PBS, centrifuge at 4°C and 300g for 3 minutes, and discard the supernatant. Repeat this step once. Resuspend the cells in 50 μL of 0.1× PBS and count number of cells with a microscope. Use the Chromium Next GEM Single Cell 3ʹ Reagent Kits v3.1 (10× Genomics, Cat No./ID: PN-1000268) for preparation of scRNA-seq library, following the manufacturer’s instructions. Libraries are sequenced in a 150-bp paired-end mode on a DNBSEQ-T7 sequencer (GenePlus, Beijing, China).

### FXcryo-Cell and cryoFX-Cell

To generate cell atlases of rice root tips, cultivated rice tiller nodes, wild rice rhizome nodes, and wounded *A. thaliana* leaves by FXcryo-Cell (Fig. 3; Figs. 4a,b; Fig. 5), we proceeded as follows: Place the plant material into ice-cold Farmers solution and apply vacuum until the material settles to the bottom of the tube. Incubate on ice for 30 minutes. Add pre-chilled 0.1× PBS to the fixed material, mix well, and incubate on ice for 5 minutes. Discard the supernatant. Repeat this step. Freeze the material in liquid nitrogen and transfer it to a −80°C freezer for long-term storage. To prepare the cells, thaw the frozen material at RT for 5 minutes and digest it with the RNase-depleted C+M+S+P digestion enzyme solution at 40°C for 30 minutes as described above. All other steps follow the FX-Cell protocol.

To generate cell atlases of the rice root tips (Fig. 3e) and the crown roots from field grown maize plants (Fig. 4d) by cryoFX-Cell, we proceeded as follows: Collect and freeze the material in liquid nitrogen and transfer it to a −80°C freezer for long-term storage. To prepare the cells, thaw the frozen material at RT for 5 minutes and digest it with the RNase-depleted C+M+S+P digestion enzyme solution at 40°C for 30 minutes as described above. All other steps follow the FX-Cell protocol.

### Genome assembly and annotation of *Oryza longistaminata*

Initial genome assembly was performed using PacBio HiFi long-read data with hifiasm (v0.19.5) under default parameters. Chromosome-scale assembly was then achieved using RagTag (v2.1.0) for scaffolding.^57^ The mitochondrial genome was assembled using oatk (v1.3.2)^58^ with specific reference files for mitochondrial and plastid assembly, employing parameters “-m angiosperm_mito.fam” and “-p angiosperm_pltd.fam”, respectively. The assembled mitochondrial sequences were aligned to the nuclear genome using minimap2 (v2.24)^59^ with parameters “--eqx -a -x asm20”. Subsequently, nuclear-mitochondrial DNA (NUMT) regions with sequence identity >= 95% were filtered out. The final genome assembly integrated the validated organellar and nuclear genomes.

Repeat annotation was conducted in two phases: First, a non-redundant transposable element (TE) library was constructed using EDTA (v2.0.0) with default plant parameters.^60^ The genome was then masked for repeats using the EDTA-derived TE library. Gene structure annotation was completed using egap (v1.0.3) with the following configurations: 1) Taxonomic constraints (taxid=4528, specific to *Oryza sativa*); 2) Protein sequence evidence from the IRGSP-1.0 reference genome (obtained from RAP-DB, https://rapdb.dna.affrc.go.jp/); 3) Tissue-specific RNA-seq data encompassing aerial (leaf/stem) and underground (root/rhizome) developmental stages of the wild-type plants. As shown in Extended Data Fig. 9, the genomes of *O. sativa* and *O. longistaminata* exhibited good synteny.

### Analysis of the scRNA-seq data generated by the 10× Genomics platform

Raw sequencing data were processed according to the manufacturer’s recommended pipeline (https://www.10xgenomics.com/cn/support/software/cell-ranger/latest). We used Cell Ranger (v7.0.0) to perform all preprocessing steps, including index building and gene expression quantification. The reference genome (TAIR10) and annotation (Araport11) for *A. thaliana* were obtained from TAIR (https://www.arabidopsis.org/). The reference genome (IRGSP-1.0) and annotation (2021-05-10) for *O. sativa* were obtained from rap-db (https://rapdb.dna.affrc.go.jp/index.html). The reference genome (B73 v5) and annotation (Zm00001eb.1) for maize were obtained from maizeGDB (https://www.maizegdb.org/). The genome assembly and annotation of *O. longistaminata* were generated in this study (see above). We implemented a comprehensive analytical workflow using Seurat v5 (https://satijalab.org/seurat/).^61^ This workflow included quality control, data normalization, dimensionality reduction, clustering, data integration, and differential expression analysis. While maintaining consistent quality control parameters within species, we applied species-specific filtering criteria across different species. Detailed quality control metrics were provided in Extended Data Table 1. Following quality control step, we performed data normalization using the “NormalizeData” function and scaled the expression values with the “ScaleData” function to enable cross-cell comparisons. Dimensionality reduction and clustering were achieved by successive applications of “RunPCA”, “RunUMAP”, “FindNeighbors”, and “FindClusters” functions. Differentially expressed genes (DEGs) were identified using the “FindAllMarkers” and “FindMarkers” functions.

### Integration of scRNA-seq datasets

To mitigate the impact of cell number discrepancies in published datasets during data integration, we employed stratified sampling for the published datasets of rice root tip^34^ and Arabidopsis leaf,^53^ retaining 51% and 38% of cells, respectively (Extended Data Table 3). For integrating the published datasets of rice root tips,^34^ maize root tips,^45^ and Arabidopsis leaves^53^ with those generated by FX-Cell, cryoFX-Cell or FXcryo-Cell, we used the “IntegrateLayers” function in Seurat.^61^ For merging the rice root tip cell atlases generated by FX-Cell, FXcryo-Cell, and cryoFX-Cell, we applied the “merge” function in Seurat.^61^ Subsequent dimensionality reduction and clustering analyses followed the aforementioned workflow.

### Cell type annotation

Cell type annotation for all single-cell datasets in this study was based on cell type marker genes derived from published literatures (Extended Data Table 1). For wild rice datasets, the orthologous genes of *O. sativa* and *A. thaliana* in wild rice were identified by BlastP, which facilitated cell type annotation through the use of established marker genes from these two species.

### GO enrichment analysis and AUCell score calculation

Gene Ontology (GO) enrichment analysis for Arabidopsis was conducted using clusterProfiler (v4.4.1)^62^ combined with the org.At.tair.db (v3.15.1) annotation package, with all parameters set to their default values. We retrieved comprehensive GO term information for the target species from the GENE ONTOLOGY database (https://geneontology.org) to construct pathway-specific gene sets. Cell-specific pathway (GO: 0009611) activity scores were then calculated using the “AUCell_run” function implemented in the AUCell package (v1.18.1),^54^ followed by downstream visualization analysis using complex heatmap.^63,64^

### Overlap coefficient analysis

To evaluate the consistency between our FX-Cell datasets and published datasets, we calculated the Szymkiewicz–Simpson coefficient (overlap coefficient)^65^ for marker gene sets identified by “FindAllMarkers” (default parameters) in the integrated rice root tip data:

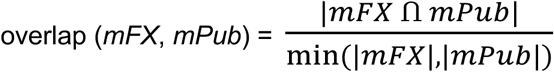

where mFX represents markers from FX-Cell data and mPub denotes markers from published data. An overlap coefficient ≥ 0.4 was considered to indicate significant similarity between gene sets.

The GO enrichment analysis was performed for each cluster’s marker genes using clusterProfiler (v4.4.1). The overlap coefficient for GO terms was calculated as:

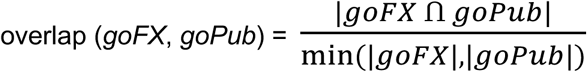

where goFX and goPub represent GO terms from FX-Cell data and published data, respectively.

### Data availability

The scRNA-seq, snRNA-seq, and RNA-seq data (BioProject PRJCA035988) were deposited in Beijing Institute of Genomics Data Center (http://bigd.big.ac.cn).^66,67^ Peer reviewers can temporally access our datasets through https://ngdc.cncb.ac.cn/gsa/s/w8ExO8OM. The chromosome-level genome assembly and annotation of *O. longistaminata* has been deposited in the Figshare database under accession number DOI 10.6084/m9.figshare.28457807.v1.

### Statistical analysis

Student’s *t*-test was performed to determine the statistical significance between different samples in quantification and phenotypes. All statistic results and graphs were generated by GraphPad Prism 8 (www.graphpad.com). The numbers of biological replicates and types of statistical analyses are given in figure legends.

